# Host preference and invasiveness of commensals in the *Lotus* and *Arabidopsis* root microbiota

**DOI:** 10.1101/2021.01.12.426357

**Authors:** Kathrin Wippel, Ke Tao, Yulong Niu, Rafal Zgadzaj, Rui Guan, Eik Dahms, Pengfan Zhang, Dorthe B. Jensen, Elke Logemann, Simona Radutoiu, Paul Schulze-Lefert, Ruben Garrido-Oter

## Abstract

Healthy plants are colonized by microorganisms from the surrounding environment, which form stable communities and provide beneficial services to the host. Culture-independent profiling of the bacterial root microbiota shows that different plant species are colonized by distinct bacterial communities, even if they share the same habitat. It is, however, not known to what extent the host actively selects these communities and whether commensal bacteria are adapted to a specific plant species. Here, we report a sequence-indexed culture collection from roots and nodules of the model legume *Lotus japonicus* that contains representatives from the majority of species found in culture-independent profiles. Using taxonomically paired synthetic communities from *L. japonicus* and the crucifer *Arabidopsis thaliana* in a multi-species gnotobiotic system, we detect clear signatures of host preference among commensal bacteria in a community context, but not in mono-associations. Sequential inoculation of either host reveals strong priority effects during the assembly of the root microbiota, where established communities are resilient to invasion by late-comers. However, we found that host preference by commensal bacteria confers a competitive advantage in their native host. We reveal that host preference is prevalent in commensal bacteria from diverse taxonomic groups and that this trait is directly linked to their invasiveness into standing root-associated communities.

---

Plant roots associate with diverse microbes that are recruited from the surrounding soil biome and which assemble into structured communities known as the root microbiota. These communities provide the host with beneficial functions, such as indirect pathogen protection, or mineral nutrient mobilization^1-3^. Despite conservation at higher taxonomic ranks^4-7^, comparison of community profiles across diverse land plants shows a clear separation according to host species^5,7^. These patterns could be explained by a process in which the root microbiota assemble according to niches defined by plant traits, that in turn diversify as a result of plant adaptation to their environment. Alternatively, variation of microbiota profiles along the host phylogeny may be at least partially caused by co-adaptation between the plant and its associated microbial communities.

Culture-independent amplicon sequencing allows characterization of community structures and taxonomic composition but does not allow the study of phenotypes of individual community members. To overcome this fundamental limitation in microbiota studies, comprehensive culture collections of sequenced strains isolated from root and leaf tissue have been established^2,3,8,9^. Synthetic communities (SynComs) built from these collections can be used in gnotobiotic reconstitution systems of reduced complexity to explore the role of immune signaling^10^, nutritional status^3,11^, biotic and abiotic stress^2^ and priority effects^12^ in the establishment of the root and leaf microbiota. However, plant microbiota studies using culture collections from different host species grown in the same natural soil have, to our knowledge, not been reported. Here, we present a bacterial culture collection from the roots and nodules of the model legume *Lotus japonicus* (hereafter *Lj*) that includes representatives of the majority of abundant species found in natural community profiles. This collection is comparable to the one previously established from *Arabidopsis thaliana* (hereafter *At*) roots^8^ in terms of taxonomic and genomic composition, despite 125 Myr of divergence between *Lj* and *At*^13^ whose crown groups evolved 65 and 32 Mya, respectively^14^.

These two collections originate from plants grown in the same soil enabling us to design ecologically meaningful SynComs for microbiota reconstitution experiments. Specifically, we allowed commensal bacteria to compete for colonization of the host from which they were derived (hereafter referred to as native) with strains isolated from the other plant species (non-native). Using this system, we explored host preference of commensal communities and the role of nitrogen-fixing nodule symbiosis, immunity and root exudation in microbiota establishment. Sequential inoculation experiments allowed us to investigate how root community assembly is affected by the order of arrival of species and to explore the link between commensal host preference and invasiveness into resident bacterial communities.

## Host species-specific bacterial culture collections

We compared the bacterial communities associated with roots of *Lj* and *At* plants grown in the same soil (Methods)^2,3,15^ and confirmed that both hosts assemble communities that are clearly distinct from those of the surrounding soil (Fig. 1a and 1b; Supplementary Data 1). This shift is characterized by a decrease in alpha-diversity (within-sample diversity; Fig. 1a) as well as by a separation between root, rhizosphere, and soil samples (beta-diversity; Fig. 1b, PCoA 2). In addition, *Lj* and *At* root samples formed two distinct clusters, indicating host species-specific recruitment of commensals from identical pools of soil-dwelling bacteria (Fig. 1b, PCoA 1), which is in line with previous studies^15,16^. This separation (28% of variance; *P* = 0.001) was mainly explained by the different relative abundance of Proteobacteria, Actinobacteria, Bacteroidetes (Flavobacteria and Sphingobacteria), and Firmicutes (Bacilli) in *Lj* compared to *At* (Supplementary Fig. 1).

**Figure 1:**
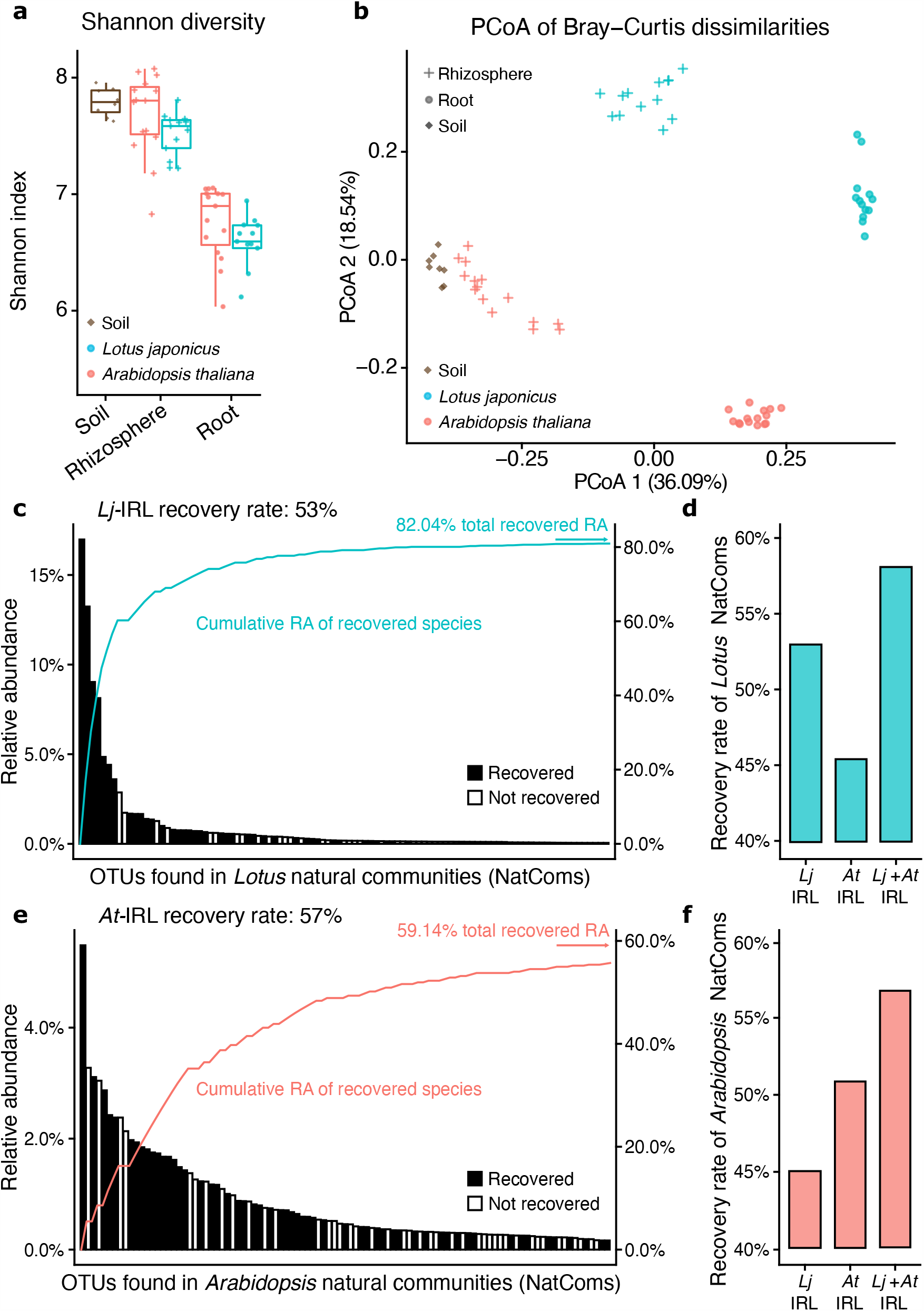
*Lotus* and *Arabidopsis* root-associated bacterial communities. **a**, Alpha-diversity analysis of soil-, rhizosphere-, and root-associated bacterial communities from *L. japonicus* and *A. thaliana* plants grown in natural soil, assessed using the Shannon index. **b**, Principal coordinates analysis (PCoA) of Bray-Curtis dissimilarities of the same communities (*n* = 64). **c** and **e**, Rank abundance plots of OTUs found in the *Lotus* (**c**) and *Arabidopsis* (**e**) natural root communities. Community members captured in the corresponding culture collection are depicted as black while non-recovered OTUs are shown in white. The vertical axis on the right shows the accumulated relative abundance in natural communities of all recovered OTUs. **d** and **f**, percentage of abundant OTUs (≥0.1% RA) associated with *Lotus-*(**d**) or *Arabidopsis* (**f**) roots in nature (natural communities, NatComs) that are captured in the *Lotus* (*Lj*-IRL) or the *Arabidopsis* (*At*-IRL) collections.

To explore the mechanisms by which different plant species assemble distinct microbial communities, we established a taxonomically and functionally diverse culture collection of the *Lj* root and nodule microbiota (Methods). A total of 3,960 colony-forming units (CFUs) were obtained and taxonomically characterized by sequencing the bacterial *16S* ribosomal RNA (rRNA; Supplementary Data 2), resulting in a comprehensive sequence-indexed rhizobacterial library from *Lj* (*Lj*-IRL). In parallel, a subset of the root samples was also subjected to amplicon sequencing to obtain culture-independent community profiles for cross-referencing with the *Lj*-IRL data. In the *Lj* collection, we were able to recover up to 53% of the most abundant bacterial OTUs (Operational Taxonomic Units, defined by 97% *16S* rRNA sequence identity) found in the corresponding natural community profiles, compared with 57% for the *At* collection (Fig. 1c and 1e; Supplementary Text and Methods). The recovered bacterial taxa in the respective collection accounted for 82% of all sequencing reads from *Lj* root samples and 59% from *At*. Approximately 45% of the abundant OTUs found in the natural communities of one host were captured in the culture collection of the other species (Fig. 1d, and 1f), indicating a substantial overlap of the recovered bacterial taxa. Both culture collections include members of the Actinobacteria, Proteobacteria, Bacteroidetes, and Firmicutes, the four phyla robustly found in the root microbiota of diverse plant species^5,7^.

To establish a core *Lj* culture collection of whole-genome sequenced strains (*Lj*-SPHERE), we selected from the *Lj*-IRL a taxonomically representative subset of bacterial isolates maximizing the number of taxa covered (Methods). A total of 294 isolates belonging to 20 families and 124 species, including both commensal and symbiotic bacteria, were subjected to whole-genome sequencing (Supplementary Data 3). Comparative analyses of all sequenced isolates from both collections revealed an extensive taxonomic and genomic overlap between exemplars derived from *Lj* and *At* (Supplementary Fig. 2; Supplementary Text). This indicates that the observed differences in natural community structures (Fig. 1b) are likely not driven by the presence of host-specific bacterial taxonomic groups. Instead, the distinct root community profiles of the two hosts are possibly due to differences in the relative abundance of shared taxonomic groups (Supplementary Fig. 1).

**Figure 2:**
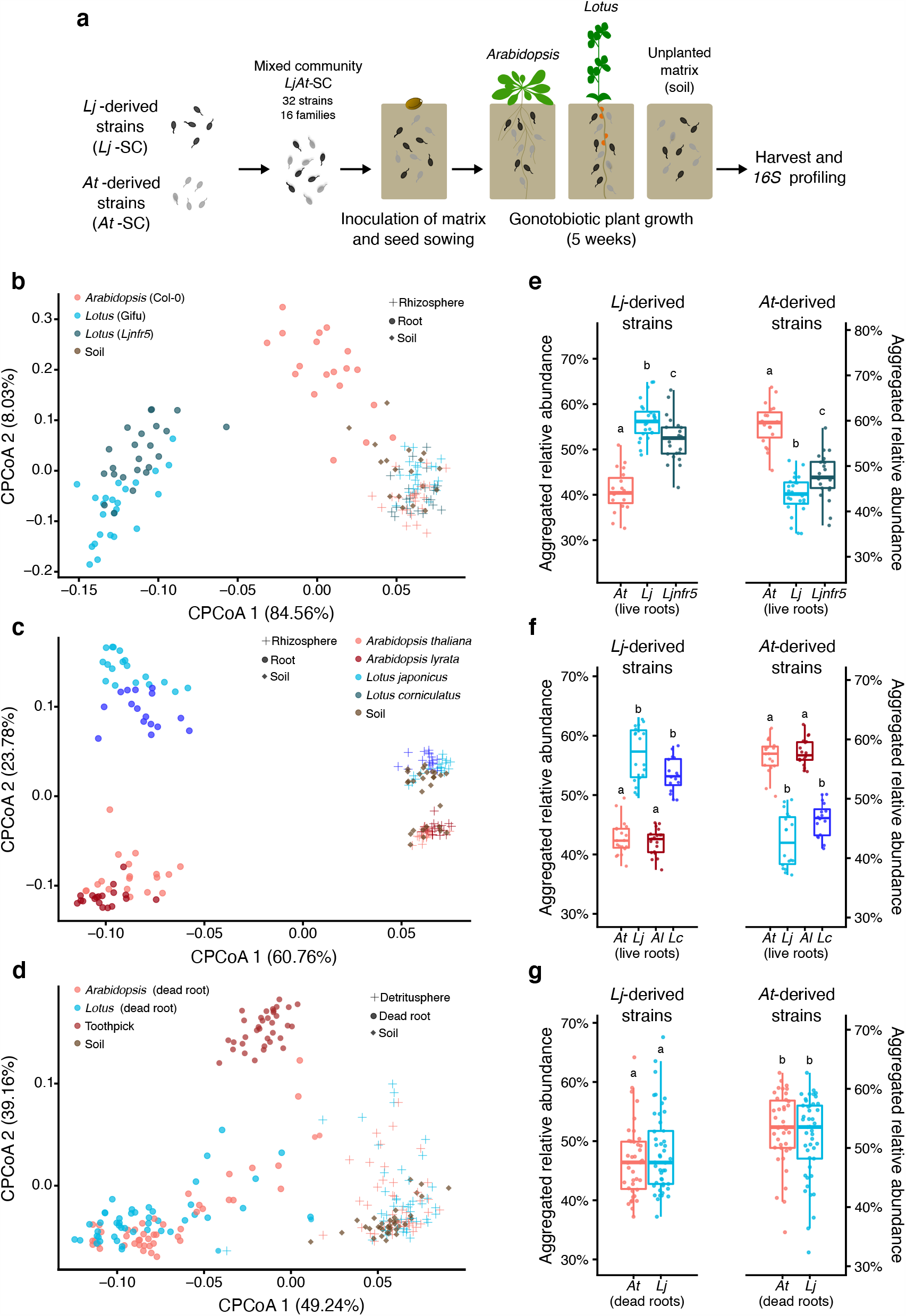
Reconstitution experiments recapitulate culture-independent patterns and show signatures of host preference by commensal communities. **a**, Setup of the competition experiments. **b, c, d**, Constrained PCoA of Bray-Curtis dissimilarities (constrained by all biological factors and conditioned by all technical variables) of soil, rhizosphere, and root samples. **b**, *L. japonicus* wild type Gifu, *nfr5* mutant, and *A. thaliana* wild type Col-0 plants co-cultivated with the mixed community *LjAt*-SC2 (experiment A, *n* = 155, variance explained 53.8%, *P* = 0.001). **c**, Gifu, Col-0, *A. lyrata* MN47, and *L. corniculatus* co-cultivated with *LjAt*-SC3 (experiment G, *n* = 173, variance explained 65.1%, *P* = 0.001). **d**, Dead roots of Gifu and Col-0, and toothpick co-cultivated with *LjAt*-SC3 (experiment H, *n* = 250, variance explained 43.9%, *P* = 0.001). **e, f, g**, Aggregated RA of the 16 *Lj*-derived and the 16 *At*-derived strains in the live (**e** and **f**) or dead roots (**g**) of *Lotus* and *Arabidopsis* plants inoculated with *LjAt*-SC2 (**e**, *n* = 66) or *LjAt*-SC3 (**f**, *n* = 72; **g**, *n* = 89).

## Host preference of commensal synthetic communities

Given the overlap between the *Lj*- and *At*-SPHERE culture collections at a high taxonomic and whole-genome level, we speculated that strain-specific phenotypic variation *in planta* could allow commensal bacteria to preferentially colonize their native host (Fig. 1b). In order to test this hypothesis, we designed taxonomically paired SynComs for each host, representing 16 bacterial families present in both collections (Supplementary Fig. 3). We then combined these SynComs into a mixed community composed of 32 strains (Supplementary Table 1) and allowed them to compete for colonization of germ-free plants. We employed a gnotobiotic system^2,17^ to grow wild-type *At* (Col-0), *Lj* (Gifu), and a *Lj* mutant deficient in root nodule symbiosis (*Ljnfr5*)^18^ in the presence of the mixed community (Fig. 2a). After five weeks, we performed community profiling *via 16S* rRNA gene amplicon sequencing of the root, rhizosphere, and unplanted soil compartments. Analysis of community diversity revealed a significant separation of communities of root samples from those of rhizosphere and soil, which in turn clustered together (Fig. 2b). In addition, we observed that the two hosts assembled distinct root microbial communities starting from the same input, and that samples from wild-type *Lj* are differentiable from those of *Ljnfr5* (Fig. 2b). These results were confirmed by an independent, full factorial experiment using a different mixed community (Supplementary Fig. 4a). An additional experiment, where strains belonging to families found exclusively in the *Lj* or *At* culture collections (two and five families, respectively) were added to the mixed community, resulted in similar patterns of beta-diversity (Supplementary Fig. 4b). These results recapitulate the community shifts between compartment, host species, and plant genotype which were previously observed in culture-independent community profiles obtained from plants grown in natural soils (Fig. 1b)^4,6,15,16^, thus validating our comparative reconstitution system to study host species-specific microbiota establishment.

**Figure 3:**
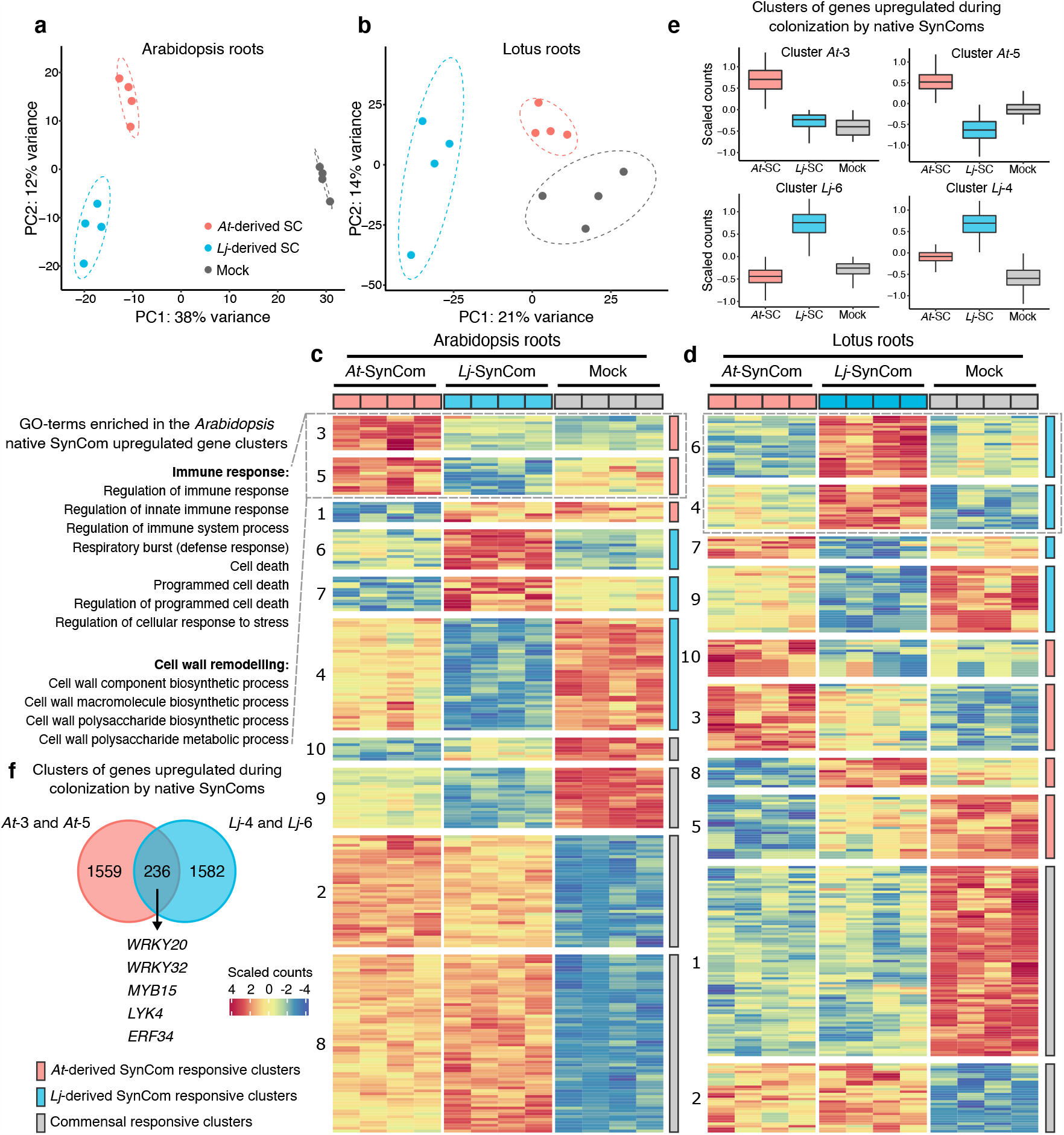
SynCom specific transcriptional outputs in Lotus and Arabidopsis roots. Whole transcriptome-level Principal Component Analysis (PC) of *Arabidopsis* (**a**) and *Lotus* (**b**) roots after co-inoculation with host-specific SynComs (*Lj*- and *At*-SC3). In the case of *Lotus* plants, a nodule isolate from the *Lj*-SPHERE collection was added to all treatments to prevent transcriptional outputs to be dominated by symbiosis or nitrogen-starvation responses. Heatmaps showing scaled counts of genes arranged according to *k*-means clustering results (only differentially expressed genes shown) for *Lotus* (**c**) and *Arabidopsis* (**d**). Distribution of expression patterns for clusters of genes upregulated after co-inoculation with native SynComs (**e**). Overlap in terms of homologues identified in the same clusters between the two host and a list of relevant transcription factors identified as potential key regulators of differential transcriptional responses (**f**).

**Figure 4:**
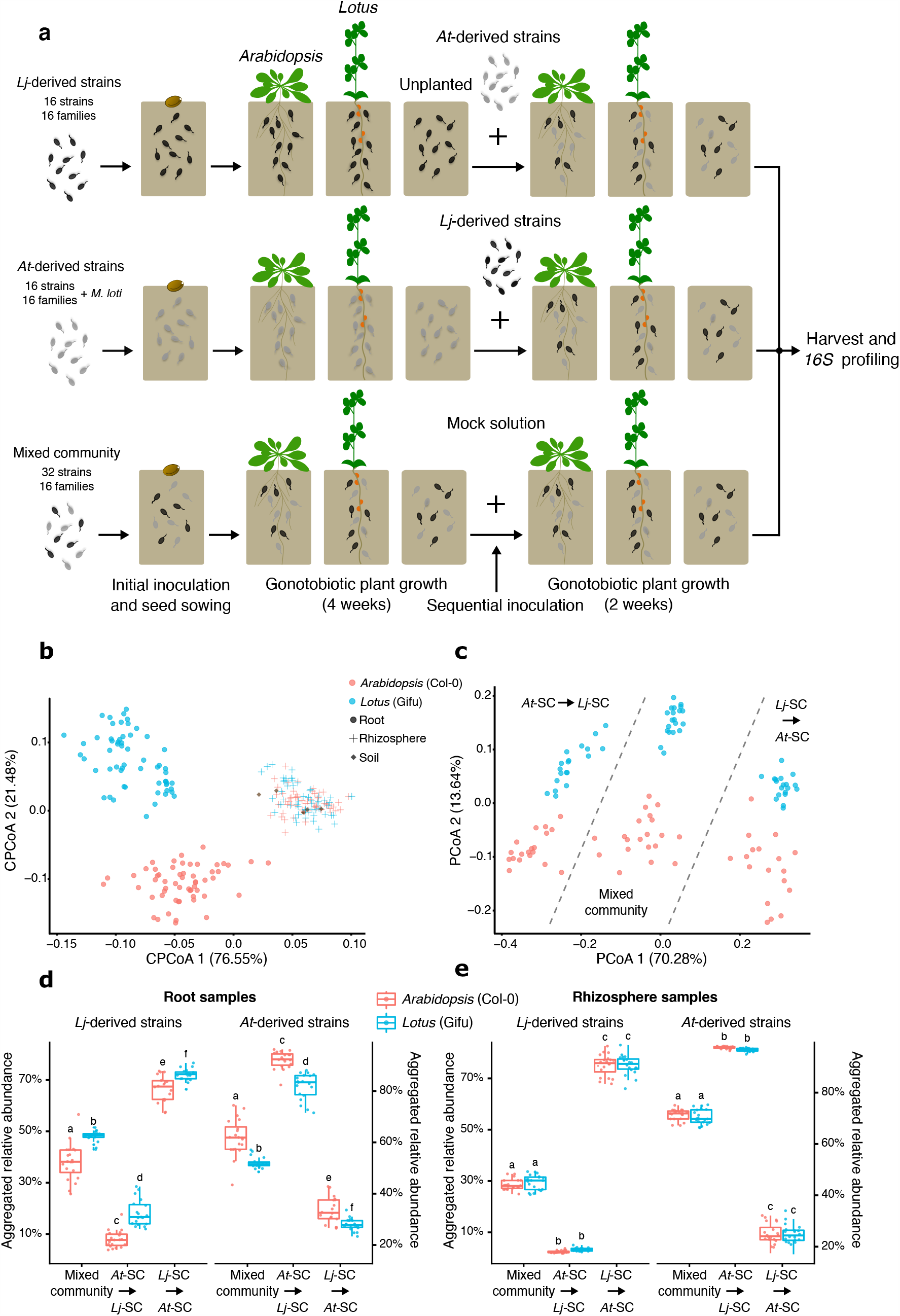
Invasion and persistence of commensal bacteria. **a**, Setup of the sequential inoculation experiment. *L. japonicus* Gifu and *A. thaliana* Col-0 plants were co-cultivated with the mixed community *LjAt*-SC3, or individual SynComs *Lj*-SC3 and *At*-SC3, followed by inoculation with the contrasting SynCom (experiment F). **b, c**, Constrained PCoA of Bray-Curtis dissimilarities (constrained by all biological factors and conditioned by all technical variables; *n* = 267; variance explained 14.7%, *P* = 0.001) of soil, rhizosphere, and root samples (**b**), and PCoA of root samples only (**c**, *n* = 137). **d, e**, Aggregated RA of the 16 *Lj*-derived and the 16 *At*-derived strains in *Lotus* and *Arabidopsis* root (**D**) (*n* = 120) and rhizosphere (**e**) (*n* = 120) samples in the indicated treatments.

Next, we tested whether communities of commensal bacteria would preferentially colonize roots of their native host species (i.e., from which they were originally isolated) compared to those of the non-native host. We found that the aggregated relative abundance of strains from the *Lj*-SPHERE collection was higher in wild-type *Lj* root samples than in those of *At* (Fig. 2e; Supplementary Fig. 4c and 4d). Likewise, strains from the *At*-SPHERE collection were more abundant on their native host than on *Lj*. Commensal host preference and host species community separation was reduced but still present in the *Ljnfr5* mutant (Fig. 2e), as well as in wild-type *Lj* after *in silico* depletion of *Mesorhizobium* reads, indicating that nodule symbiosis only partially contributes to commensal host preference. Further, sequential *in silico* removal of individual bacterial families did not significantly alter the observed patterns of host preference at the community level (Supplementary Fig. 5). Mono-association experiments with *Lj* and *At* wild-type plants grown on agar plates revealed that most community members did not show a significant host preference in isolation (Supplementary Fig. 6), suggesting that this commensal phenotype requires a community context. Moreover, we found that shoot biomass of both host species was not negatively affected by these strains, confirming their commensal lifestyle in mono-associations (Supplementary Fig. 7).

**Figure 5:**
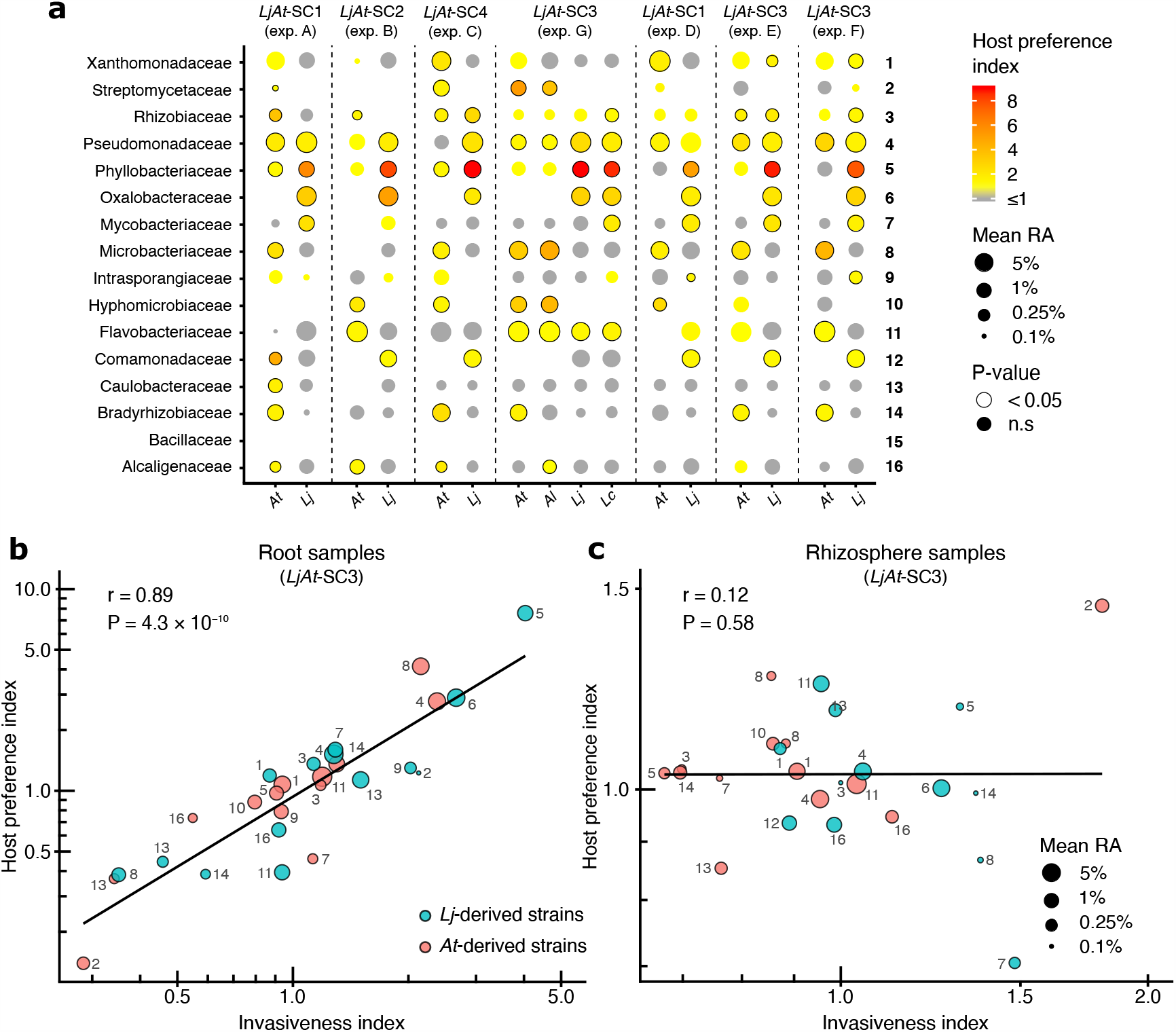
Host preference is linked to invasiveness. **a**, Analysis of host preference of individual commensal strains across gnotobiotic experiments (*n* = 366). Each strain is represented by a dot, whose color corresponds to its host preference index and whose size to its average relative abundance. A significant host preference (Mann-Whitney test, FDR-corrected) is depicted by a black circle around a dot. **b, c**, Correlation between host preference and invasiveness index for each strain in root (*n* = 115) and rhizosphere samples (*n* = 119), respectively, obtained from the sequential inoculation experiment (exp. F). The color of each point designates the host of origin of each strain and the size denotes its mean relative abundance (log2-transformed). Each point is labeled with a numeric identifier that corresponds to the strains in panel **a** (*LjAt*-SC3).

We then investigated if the phenotype of commensal host preference was conserved in a plant phylogenetic framework. We selected two additional plant species, *L. corniculatus* and *A. lyrata*, which diverged from *Lj* and *At* approximately 12.5 Mya and 13 Mya ago, respectively^19,20^, and are endemic to the region from which the soil used to isolate these bacterial strains was collected^21,22^. We inoculated these four species with a mixed community of *Lj* and *At* commensals and obtained amplicon profiles of root, rhizosphere and unplanted soil samples. We observed a significant separation between *Lotus* and *Arabidopsis* root communities (Fig. 2c; *P*=0.001), and to a lesser extent between samples from the sister species within the same genus (Supplementary Fig. 8), which is in line with similar results obtained from *At* relatives grown in natural sites^23^. We found that the patterns of host preference observed in *Lj* and *At* were retained in their relative species (Fig. 2f), suggesting that this community phenotype might be the result of commensal adaptation to root features conserved in a given host lineage.

## Host factors driving preferred associations in the root microbiota

Previous studies have reported shifts in *At* leaf or root microbiota structure in mutants impaired in different host immunity pathways^10,24^. We speculated that the plant immune system might also play a role in selecting commensal bacteria in a host-specific manner. We thus tested whether host mutants impaired in perception of ubiquitous microbe-associated molecular patterns (MAMPs) were also preferentially colonized by native commensal strains. Community profiles of roots of *At* and *Lj* mutants lacking the receptor FLS2, which detects the bacterial flagellin epitope flg22 (*Ljfls2* and *Atfls2*)^25,26^, were indistinguishable from those of their respective wild types (Supplementary Fig. 9a). Similar results were obtained with an *At* mutant lacking MAMP co-receptors BAK1 and BKK1 as well as CERK1 receptor kinase, known to play a role in the perception of the bacterial MAMP peptidoglycan (*Atbbc* triple mutant)^27^. In addition, bacterial host preference was retained in those mutants (Supplementary Fig. 9b). A separate experiment using the *dde2 ein2 pad4 sid2* (*deps*) mutant in *At*, which is simultaneously defective in all three major defense phytohormone signaling pathways (salicylic acid, jasmonate and ethylene)^28^, showed comparable results (Supplementary Fig. 9c and 9d). Together, these data suggest that the tested MAMP receptors and immune signaling pathways do not play a crucial role in preferential colonization by native commensal bacteria.

Plant root exudates contain molecular cues that can be differentially metabolized or perceived as signals by root microbiota members^29,30^. In particular, glucosinolates (GS), a group of nitrogen- and sulfur-containing metabolites found in root exudates throughout the family Brassicaceae, including *At*, are known to play a role in plant defense and serve as precursor of compounds that inhibit microbial growth^31-33^. Since legumes such as *Lj* lack genes required for GS biosynthesis, we speculated that secretion of these compounds by *At* might contribute to the observed differences in community structure. We therefore tested whether the *At cyp79b2 cyp79b3* double mutant^34^, which is defective in the production of microbe-inducible and tryptophan-derived metabolites, including indole GSs, was also preferentially colonized by native commensal strains. Comparison of bacterial community profiling data suggests that indole GS had no impact on overall community structure or bacterial host preference *in planta* (Supplementary Fig. 10). Notably, incubation of bacterial SynComs in root exudates from *Lj* and *At* plants in an *in vitro* millifluidics system (Methods) resulted in small but significant community separation according to the plant genotype (Supplementary Fig. 11a; 5% of variance; *P*=0.002). However, in this system, we observed a loss of the host preference phenotype (Supplementary Fig. 11b), which prompted the question of whether live root tissue was required for preferential colonization by native commensals. We profiled the bacterial communities associated with dead root material from flowering *Lj* and *At* wild-type plants and with inert lignocellulose matrices (softwood birch toothpicks) at 5, 12, and 19 days after inoculation with a mixed community. Diversity analyses showed that dead roots and toothpicks harbored distinct microbial communities that were separated from those of soil or detritusphere (soil surrounding dead roots), independently of the timepoint (Fig. 2f). This separation was likely driven by a significant increase in the abundance of Flavobacteria, a taxon associated with the capacity to decompose complex polysaccharides^35^, and which dominates the dead root communities (53% RA on average). Unlike the large separation between living *Lj* and *At* roots (36% of the variance), we observed only a small but significant differentiation between *Lj* and *At* dead root communities (6.4% of variance; *P*=0.001). Additionally, commensal host preference was undetectable in dead roots, where *Lj*- and *At*-derived strains reached similar aggregated relative abundances in root material harvested from either host (Fig. 2g). Taken together, these results suggest that a living root and other factors besides root exudates, such as a physical contact with the plant (i.e., host-commensal feedbacks) are required for host preference in the root microbiota.

## SynCom-specific transcriptional responses of *Lj* and *At* roots

Next, we sought to assess whether native, non-native or mixed commensal communities elicited a differential response in either host species. We grew wild-type *Lj* and *At* plants in our soil-based gnotobiotic system inoculated with *Lj*-, *At*- or mixed SynComs for five weeks. Assessment of plant performance revealed that treatment with commensal communities leads to increased plant biomass compared to axenic controls. However, no significant differences in plant biomass were observed between the different SynCom treatments (Supplementary Fig. 12). Next, we conducted RNA sequencing of roots and found that transcriptional outputs separated according to SynCom treatment in both hosts (Fig. 3). Analysis of *k*-means clustering of whole transcriptomes revealed gene clusters associated with general response to bacterial colonization, as well as clusters specific to treatment with native or non-native SynComs. Among genes specifically induced by the native SynComs we found several transcriptional regulators of immunity (e.g. WRKY20, WRKY32, and MYB15), well characterized MAMP receptor kinases (LYK4) and ethylene response factors (e.g. ERF34). This conserved pattern of differential response in the two plant species suggests a specific transcriptional response to native commensal communities that involves components of the host immune system. The differentially expressed transcription factors identified here constitute prime candidates for future exploration of the underlying mechanisms of host preference.

## Invasiveness and persistence in the root microbiota

The results obtained from four independent experiments using four different mixed communities (Fig. 2; Supplementary Figs. 4, 9 and 10) show that native strains have a competitive advantage when colonizing roots of their cognate host. Ecological theory suggests that in the presence of a competitive hierarchy, the order of species arrival does not matter, as better adapted species tend to dominate irrespective of the history of the community^36^. To investigate the role that priority effects play in root community assembly we designed a series of sequential inoculation experiments using host-specific SynComs (Fig. 4a; Supplementary Table 2 and Supplementary Data 4). *At* and *Lj* wild-type plants were inoculated with taxonomically paired SynComs derived from *Lj* (*Lj*-SC3), *At* (*At*-SC3) or a mixed community (*LjAt*-SC3) for four weeks. Subsequently, we challenged the established root communities by adding the complementary SynCom (*At*-SC3 or *Lj*-SC3, respectively) to the soil matrix or, in the case of plants initially treated with the mixed community (*LjAt*-SC3), a mock solution (Fig. 4a). We then allowed all plants to grow for an additional two weeks before harvesting. Amplicon sequencing showed a significant separation of communities by compartment, and, within root samples, according to host species (Fig. 4b; *P*=0.001), mirroring the patterns observed in natural communities (Fig. 1a). Interestingly, analysis of beta-diversity of root samples at strain-level resolution revealed an effect of the treatment on community structure (Fig. 4c), demonstrating that the order of arrival of strains affects community outputs. Examination of aggregated relative abundances showed that, in a competition context (i.e., initial inoculation with the mixed community *LjAt*-SC3), commensal SynComs preferentially colonized roots of their cognate host (Fig. 4d), in line with results from the previous competition experiment shown in Figure 2. However, in an invasion context, early-arriving SynComs invariably reached higher proportions in the output communities compared to the late-arriving SynComs (Fig. 4d and 4e). Notably, estimation of absolute bacterial abundances showed that a secondary inoculation with an invading SynCom did not result in a significant increase in total bacterial load (Supplementary Fig. 13). Together, the results from our sequential inoculation experiments (Fig. 4C and D) are indicative of the existence of priority effects in the root microbiota. These effects could be explained by niche preemption, where early-arriving community members reduce the amount of resources available (e.g. nutrients, space) for latecomers^37^; alternatively, they could be the result of a feedback process between the host and the early-arriving commensals.

We hypothesized that commensal bacteria would be less affected by priority effects when colonizing their cognate host, given their competitive advantage with respect to non-native strains. In order to test this, we examined aggregated relative abundances of *Lj*- and *At*-derived SynComs in the root and rhizosphere communities. We found that host-specific SynComs were better able to invade a resident community in the roots of their native host compared to those of the other plant species (Fig. 4d), thus reducing the strength of the priority effects. However, in the rhizosphere compartment of either plant species, host-specific SynComs showed neither host preference in a competition context nor differences in their ability to invade standing communities (Fig. 4e).

We then tested if host preference was directly linked to invasiveness and to what extent these traits were found in individual community members. First, we quantified the strength of host preference by calculating the ratio between the relative abundance of each strain in their native host compared to the other plant species (host preference index; Methods). Notably, although *Lj* root samples did not include nodules, but possibly contained incipient symbiotic events, the strains with the highest host preference index were the nitrogen-fixing *Lj* symbionts belonging to the Phyllobacteriaceae family (Fig. 5a), indicating that host preference of symbiotic rhizobia is not limited to nodule tissue. In addition, multiple other commensal strains showed significant host preference, with members of the families Pseudomonadaceae, Rhizobiaceae, Oxalobacteriaceae, and Comamonadaceae robustly displaying a high host preference index. Next, we calculated an invasiveness index by comparing the ability of each strain to invade a standing community on their native host compared to the other plant species (Methods). We found a strong correlation between host preference and invasiveness of commensal bacteria which is independent of their relative abundance (*r* = 0.89; *P* = 4.3×10^−10^; Fig. 5b). In contrast, this correlation was absent in the rhizosphere samples (Fig. 5c), indicating that the link between these two bacterial traits is mediated by host attributes that do not extend to the rhizosphere. Together, our data show that host preference is prevalent in commensal bacteria from diverse taxonomic groups and that this trait is tightly linked to invasiveness and together play a role during root microbiota assembly.

## Conclusions

The current concept of host specificity in plant-microbe interactions was originally developed based on studies using microorganisms with either pathogenic or mutualistic lifestyles. Recently, it has been shown that soilborne, nitrogen-fixing *Ensifer meliloti* symbionts can adapt to local host genotypes in only five plant generations, and proliferate to greater abundances in hosts with shared evolutionary histories^38^. We show here that in the *Lj* and *At* root microbiota, there is a gradient of host preference among commensals belonging to diverse taxonomic lineages. Maintenance of host preference in the sympatric relative species *L. corniculatus* and *A. lyrata* raises the possibility that these commensals might have adapted to host features conserved in the respective plant genera. Alternatively, the observed host preference patterns might be the consequence of other ecological processes, such as ecological fitting, whereby organisms are able to colonize and persist in a new environment using traits that they already possess^39^. Diversification of plant traits as a result of adaptation to edaphic or other environmental factors is expected to result in new host features that constitute novel root niches for microbial colonization. It is also possible that host diversification is partly driven by the adaptation of plants to commensal microbiota in soils with contrasting properties. The physiological relevance for the plant might be explained by the observation that similarity between the root microbiota of different species affects competitive plant-plant interactions and has an impact on host performance through plant-soil feedback^5^.

In aquatic and terrestrial ecosystems, microbial traits such as growth rate, antagonistic activity or resource use efficiency are known determinants of invasiveness^40,41^. In microbial communities associated with a eukaryotic organism, the ability to interact with the host might also be required for successful invasion. Our results indicate that native commensals have a competitive advantage when invading standing communities in the root but not in soil or rhizosphere. One possibility is that increased invasiveness by native bacteria is enabled by the existence of unfilled host species-specific root niches that can be occupied by late-comers. Alternatively, direct interaction of commensals with their host may be required to trigger the formation of host species-specific root niches, which could be linked to the specific transcriptional reprogramming in roots observed during colonization by native SynComs. This latter hypothesis is further supported by the observation that bacterial SynComs colonizing dead roots or incubated in root exudates *in vitro* showed no significant host preference. Our study provides a framework to test these hypotheses by exploring, in future experiments, the molecular basis of host preference in multiple taxa of the bacterial root microbiota in comparison with host adaptation mechanisms in plant pathogens and symbionts.

## Methods

### Bacterial and plant material and growth conditions

Most bacterial strains were grown in tryptic soy broth (15 g/L, TSB, Sigma-Aldrich) liquid medium or on agar plates containing 15 g/L of Bacto Agar (Difco) at 25°C. *Mesorhizobium* strains LjNodule210 and LjNodule218, isolated from *Lotus japonicus* wild-type root nodules, were cultured in TY medium (5 g/L tryptone, 3 g/L yeast extract) supplemented with 10 mM CaCl_2_ or in YMB medium (5 g/L mannitol, 0.5 g/L yeast extract, 0.5 g/L K_2_HPO_4_·3H_2_O, 0.2 g/L MgSO_4_·7H_2_O, 0.1 g/L NaCl). The composition of synthetic bacterial communities (SynComs) is listed in Supplementary Table 1. The *L. japonicus* ecotype Gifu B-129 was used as wild type, and the symbiosis-deficient mutant *nfr5-2*^18^, and flagellin receptor-deficient mutant *fls2* (LORE1-30003492)^25^ were derived from the Gifu B-129 genotype. For *Arabidopsis thaliana*, ecotype Columbia-0 was used as wild type. The mutant genotypes *fls2*^26^, *bbc*^27^, *deps*^28^, and *cyp79b2 cyb79b3*^34^ were available in our seed stock. *L. corniculatus* seeds, cultivated in the North-Western German lowland, were retrieved from Rieger-Hofmann GmbH, Blaufelden-Raboldshausen, Germany. *A. lyrata* MN47 seeds were a gift from Prof. Juliette de Meaux, University of Cologne.

### Establishment of the *L. japonicus* root-associated bacterial culture collection

The *L. japonicus* culture collection combines strains isolated during three independent isolation events. Bacterial isolation, DNA isolation, and identification using Illumina sequencing were essentially performed as previously described^8^. Wild-type *L. japonicus* (ecotype Gifu B-129) plants were grown in natural soil (Cologne agriculture soil, CAS, batch 10 from Spring 2014, and batch 11 from spring 2015) in the greenhouse and harvested after four or eight weeks to cover different developmental stages. Root systems of 20 plants were subjected to DNA isolation and culture-independent community profiling *via* amplicon sequencing (see below). From 45 plants, a 4-cm section of the roots was collected and rigorously washed three times with phosphate-buffered saline (PBS; 130 mM NaCl [7.6 g/l], 7 mM Na_2_HPO_4_ [1.246 g/l], 3 mM NaH_2_PO_4_ [0.414 g/l], pH 7.0) and three times with sterile water. Nodule and root parts were separated and homogenized independently. Homogenized roots from each individual plant were allowed to sediment for 15 min and the supernatant was diluted (1:20K, 1:40K, and 1:60K) with four different media: 3 g/L TSB, CY, 50%TY, and YAN, a medium enriching for Burkholderiales (10 g/L yeast extract, 1 g/L K_2_HPO_4_, and 0.5 g/L MgSO_4_·7H_2_O). Bacterial dilutions were distributed and cultivated in 96-well microtiter plates. Homogenized nodules from each individual plant were directly diluted (1:20K, 1:40K and 1:60K), distributed and cultivated in 96-well microtiter plates. This procedure was carried out for individual plants to obtain bacterial isolates from different plant roots to capture the intra-species genetic diversity of the bacterial isolates. After 10–20 days of incubation at room temperature, plates that showed visible bacterial growth in around 30 wells were chosen for high-throughput Illumina sequencing. For identification of the bacterial isolates, a two-step barcoded PCR protocol described previously^8^ was utilized, with the difference that at the first step of the PCR, the v5-v7 fragments of the *16S* rRNA gene were amplified by the degenerate primers 799F (AACMGGATTAGATACCCKG) and 1192R (ACGTCATCCCCACCTTCC), and indexing was done using Illumina-barcoded primers. The indexed *16S* rRNA amplicons were subsequently pooled, purified, and sequenced on the Illumina MiSeq platform. Strains isolated from nodules were subsequently tested for their ability to form functional nodules in *L. japonicus* Gifu plants grown on agar plates.

Next, cross-referencing of IRL sequences with culture-independent profiles allowed us to identify candidate strains for further characterization, purification, and whole-genome sequencing. Two main criteria were used for this selection: first, we aimed at obtaining maximum taxonomic coverage and selected candidates from as many taxa as possible; second, we gave priority to strains whose *16S* sequences were highly abundant in the natural communities. Whenever multiple candidates from the same phylogroup were identified, we aimed at obtaining multiple independent strains, if possible, coming from separate biological replicates to ensure they represented independent isolation events. After validation of selected strains, 294 were successfully subjected to whole-genome sequencing.

For whole-genome sequencing, DNA was isolated from strains using the QiAmp Micro DNA kit (Qiagen, Hilden, Germany), treated with RNase, and purified. Quality control, library preparation, and sequencing (on the Illumina HiSeq3000 platform) were performed by the Max Planck-Genome center, Cologne, Germany (https://mpgc.mpipz.mpg.de/home/).

Sequencing depth was 5 million reads per sample.

### Culture-independent community profiling of *Lotus* and *Arabidopsis* root and corresponding soil samples

Bacterial communities were profiled by amplicon sequencing of the variable v5-v7 regions of the bacterial *16S* rRNA gene. Library preparation for Illumina MiSeq sequencing was performed as described previously^2^. In all experiments, multiplexing of samples was performed by double-indexing (barcoded forward and reverse oligonucleotides for *16S* rRNA gene amplification).

### Greenhouse experiment

*L. japonicus* Gifu and *A. thaliana* Col-0 were grown for five weeks in CAS soil (batch 15 from January 2020) in 7×7 cm pots alongside unplanted control pots under short-day conditions. Pots were watered with sterile water from the bottom as needed. Root, rhizosphere, and soil samples were harvested and processed as described previously^42^. In total, 15, 13, and 8 replicates were sampled for Col-0, Gifu, and unplanted controls, respectively. DNA was isolated from those samples using the MP Biomedicals FastDNATM Spin Kit for Soil.

### Multi-species microbiota reconstitution experiments

We utilized the gnotobiotic FlowPot system^2,17^ to grow *A. thaliana* and *L. japonicus* plants with and without bacterial SynComs. In brief, the system allows for even inoculation of each growth pot with microbes by the flushing of pots with the help of a syringe attached to the bottom opening. Subsequently, sterilized seeds are placed on the matrix (peat and vermiculite, 2:1 ratio), and pots are incubated under short-day conditions (10 hours light, 21°C; 14 hours dark, 19°C), standing in customized metal racks in sterile plastic boxes with filter lids (SacO2 microboxes, www.saco2.com). For SynCom preparation, bacterial commensals were grown separately in liquid culture for 2-5 days to reach high density, harvested, and washed in 10 mM MgSO_4_. Equivalent amounts of each strain were combined to yield the desired SynComs with an optical density (OD_600_) of 1. An aliquot of 200 µL of the SynCom as reference sample for the experiment start, and aliquots of 50 µL of the individual strains were taken and stored at −80°C for sequencing. The SynCom was added to the desired medium to reach a final OD_600_ of 0.02. FlowPots were each flushed with 50 mL of inoculum (medium/SynCom mix). Generally, the medium used for inoculation was 0.25x B&D^43^ supplemented with 1 mM KNO_3_ for both plant species. In experiments C, G, and J (Supplementary Table 2), 0.5x MS (2.22 g/L Murashige+Skoog basal salts, Duchefa; 0.5 g/L MES anhydrous, BioChemica; adjusted to pH 5.7 with KOH) was used for *Arabidopsis*. The two plant species were grown in separate FlowPots side-by-side, with ten pots in total per plastic box. After five weeks of growth, roots were harvested and cleaned thoroughly from attached soil using sterile water and forceps. Root segments of *Lotus* containing root nodules were omitted from sampling. Soil samples from planted pots were collected as rhizosphere samples, and soil samples from unplanted pots were collected as soil samples. All root (comprising both the epiphytic and endophytic compartments), rhizosphere (soil from planted pots), and soil (soil from unplanted pots) samples were transferred to Lysing Matrix E tubes (FastDNA™ Spin Kit for Soil, MP Biomedicals), frozen in liquid nitrogen, and stored at −80°C for further processing. DNA was isolated from those samples using the MP Biomedicals FastDNA™ Spin Kit for Soil, and from individual strains of the SynCom *via* quick alkaline lysis^8^, and subjected to bacterial community profiling or absolute quantification of bacteria. When RNA was isolated, samples were harvested the same way and processed using the RNeasy Plant Mini kit (Qiagen, Hilden, Germany).

### Dead root experiment

*L. japonicus* Gifu and *A. thaliana* Col-0 plants were grown in potting soil in the greenhouse.Mature root systems were harvested from flowering plants (13-week old *Lotus*, 7-week old *Arabidopsis*), washed several times in water, padded on kitchen paper to remove moisture, and dried in big glass petri dishes at 120°C for one hour. Note that Gifu plants had a few small, most likely ineffective root nodules. Pieces of the dried, dead roots were planted into FlowPots under sterile conditions, and inoculation with a SynCom (*LjAt*-SC3) was performed as described above. Dead roots were recovered from the FlowPots after 5, 12, and 19 days of incubation, and washed and stored as described above for live roots.

### SynCom invasion experiments

To test the capacity of native and non-native commensals to invade a standing root-associated community, we sequentially inoculated FlowPots with native and non-native strains. FlowPots were prepared as usual, with the addition of a round nylon filter (pore size 200 µm) at the bottom of the pot to avoid clogging of the bottom opening by matrix material. FlowPots were first inoculated with either the mixed SynCom (32 strains, derived from both *A. thaliana* and *L. japonicus*), the *At* SynCom (16 strains derived from *A. thaliana*), the *Lj* SynCom (16 strains derived from *L. japonicus*), or the mock solution (medium only). The medium used for inoculation was 0.25x B&D^43^ supplemented with 1 mM KNO_3_ for both plant species.

Sterilized *A. thaliana* Col-0 seeds and germinated sterile *L. japonicus* Gifu seeds were placed on the soil surface. Note that a few drops of *Mesorhizobium* culture (*Lotus* root nodule symbiont, strain LjNodule218, OD_600_ 0.02) were applied to Gifu seedlings in the *At* SynCom treatment to allow for normal root nodule symbiosis to occur and ensure healthy plant growth. After growth of Col-0 and Gifu plants for four weeks, a second inoculation was performed, where a mock inoculum (medium) was added to the mixed SynCom-treated pots, the *Lj* SynCom was added to the *At* SynCom-treated pots, the *At* SynCom was added to the *Lj* SynCom-treated pots, and mock inoculum was added to the mock-treated pots. Because the routine flushing with a syringe through the bottom was not possible while plants were growing in the pots, we flushed the pots in reverse by adding the inoculum from the top and applying vacuum from the bottom. On a sterile bench, FlowPots (which essentially are cut 60-mL syringes with a male Luer Lok connector) were placed onto female Luer Lok connectors of a vacuum manifold (QIAvac 24 Plus, Qiagen), keeping the valves of the manifold closed. Vacuum was applied to the manifold with an attached vacuum pump. Then, 20 mL of inoculum were carefully added to one pot with a 20-mL syringe and needle, making sure not to damage the plant shoots. The valve where this pot was attached to was opened, the liquid sucked through the FlowPot, and the valve closed again. This was repeated until all pots had been inoculated. Pots were put back into the plastic containers and plants were grown for another two weeks. Root, rhizosphere, and soil samples were harvested as described above.

### Collection of root exudates

*Arabidopsis* and *Lotus* plants were grown in a customized hydroponic system (original design by Manuela Peukert, University of Cologne, unpublished). This sterile growth setup consists of glass jars filled with glass beads and a stainless-steel mesh on top. Nutrient solution (modified 0.25x B&D medium; Fe-EDTA instead of Fe-citrate) was poured into the jars until the beads were covered in liquid and the liquid touched the metal mesh. We employed the same medium for both plant species in order to allow for direct comparison of exudate composition, and to minimize differential effects on the bacterial community originating from different media types. We chose the *Lotus* B&D medium since *Arabidopsis* grew reasonably well in it. Sterilized and pregerminated seeds were placed onto the mesh, jars were put into sterile plastic boxes with filter lids (SacO2 microboxes), and plants were grown for five weeks. The medium containing root exudates was removed from the jars in the clean bench using a sterile metal needle and plastic syringe. After transfer to 50-mL Falcon tubes, exudates were frozen at −80°C, freeze-dried until a volume of 2-3 mL was left, thawed, and adjusted with sterile water to 5 mL. Exudates were kept at −80°C until further usage.

### Millifluidics experiment

Incubation of commensal strains in root exudates was performed in a novel millifluidics system (MilliDrop Analyzer, MilliDrop, Paris, www.millidrop.com). This drop-based system allows us to incubate bacteria in very small volumes of root exudates or growth medium. In brief, bacteria and exudates or growth medium are combined in wells of a 96-well plate using a pipetting robot Freedom Evo 100® (Tecan, France). Droplets of approximately 100-200 nL in volume are then sucked in from the wells of the loading plate by a tip on the robotic arm of the MilliDrop Analyzer, generating hundreds of droplets within an oil-filled tube, separated by air spacers. During incubation, the droplet “train” moves back and forth, so that during each round, each droplet passes a detector that counts the droplets and records fluorescence. The growth of microbial organisms in this system can be monitored by adding the vital dye resazurin to the cultures, which is reduced to red-fluorescent resorufin by microbial metabolic activity. Culture droplets can be collected after the experiment and subjected to community profiling.

For this experiment, the mixed community *LjAt*-SC1 (Supplementary Table 1) was used and was essentially prepared as described above for the *in planta* experiments. It was adjusted to OD_600_ of 0.1 and used as input for preparation of the loading plate. Pure exudates (pH between 7.0 and 8.0) or a defined M9+carbon growth medium (1x M9 salts including phosphate buffer, 1 mM magnesium sulfate, 0.3 mM calcium chloride, 1x vitamin B solution, and artificial root exudates, pH 7.0) was used for incubation. Vitamin B solution contained 0.4 mg/L 4-aminobenzoic acid, 1 mg/L nicotinic acid, 0.5 mg/L calcium-D-pantothenate, 1.5 mg/L pyridoxine hydrochloride, 1 mg/L thiamine hydrochloride, 0.1 mg/L biotin, and 0.1 mg/L folic acid (modified from Pfennig, 1978^44^). Artificial root exudates (modified from Baudoin *et al*. 2003^45^) were composed of 0.9 mM glucose, 0.9 mM fructose, 0.2 mM sucrose, 0.8 mM succinic acid, 0.6 mM sodium lactate, 0.3 mM citric acid, 0.9 mM serine, 0.9 mM alanine, and 0.5 mM glutamic acid. Bacteria were incubated for three days, during which the pH of the cultures stayed stable. Droplets were collected in 6 µL, and DNA isolated via quick alkaline lysis^8^, which consisted of addition of 10 µL of buffer 1 (25 mM NaOH, 0.2 mM EDTA, pH 12), incubation at 95°C for 30 min, addition of 10 µL of buffer 2 (40 mM Tris-HCl at pH 7.5), storage at −20°C.

### Mono-associations of SynCom members with host plants

*Lotus* seeds were sterilized and placed on sterile wet Whatman paper for germination. Seedlings were transferred to squared petri dishes containing 0.25x B&D medium (with Fe-EDTA instead of Fe-citrate) supplemented with 3 mM KNO_3_ and 1% Difco Bacteriological agar, and sterile filter paper was put on top of the sloped solidified medium prior to placing the seedlings to prevent root growth inside the agar. *Arabidopsis* seeds were sterilized and germinated on 0.5x MS medium plus 1% Difco Bacteriological agar. Seedlings were transferred to squared petri dishes containing 0.5x MS medium (neutral pH, buffered with 2 mM HEPES) plus 1% agar. The 32 strains of the mixed community *LjAt*-SC3 were grown individually in liquid medium, harvested, and adjusted to an OD_600_ of 0.02. Seedlings were inoculated by adding 500 µL of bacterial culture to the roots. Plants were grown for 14 days under long-day conditions (16/8 day-night cycles) at 21°C. Three biological replicates were prepared for each genotype-bacteria combination.

### Absolute quantification of bacteria in root samples via qPCR and colony counting

Genomic DNA was isolated from roots of plants grown in FlowPots for six weeks (experiment F, Supplementary Table 2), using the MP Biomedicals FastDNA™ Spin Kit for Soil. DNA concentration was determined fluorometrically using the Quant-iT PicoGreen dsDNA Assay Kit (Thermo Fisher Scientific). To quantify bacterial load on plant roots, the amount of bacterial DNA relative to the amount of plant DNA was determined *via* qPCR. For bacteria, the v5-v7 region of the *16S* rRNA gene was amplified using the AACMGGATTAGATACCCKG (799F) and ACGTCATCCCCACCTTCC (1192R) primers.

For *A. thaliana* Col-0, a fragment of At1g12360 was amplified using the TCCGGTCAATATTTTTGTTCG and TATAGCAGCGAAAGCCTCGT primers, and for *L. japonicus* Gifu, a fragment of the *NFR5* gene was amplified using the TCATATGATGGAGGAGTTGTCTGTT and ATATGAGCTTCGGAGCATGG primers.qPCR was performed as described previously^46^. The amount of *16S* rRNA was normalized to plant gene within each individual sample using the following equation: *16S* rRNA gene over plant gene = 2^-Ct(16S)^ / 2^-Ct(plant)^.

For colony counts, roots were harvested, washed, weighed, and crushed in 500 µl (Col-0) or 750 µl (Gifu) sterile water. Serial dilutions of 10^−1^, 10^−2^, 10^−3^, 10^−4^, and 10^−5^ of the crushed roots were prepared in sterile water. 10 µl each were spotted onto 10% TSB agar square plates. Single colonies were counted after 1-3 days.

### Processing of *16S* rRNA gene amplicon data

Amplicon sequencing data from *L. japonicus*^15^ and *A. thaliana*^2^ roots of plants grown in CAS soil in the greenhouse, along with unplanted controls, were demultiplexed according to their barcode sequence using the QIIME^47^ pipeline. Afterwards, DADA2^48^ was used to process the raw sequencing reads of each sample. Unique amplicon variants (ASVs) were then inferred from error-corrected reads, followed by chimera filtering, also using the DADA2 pipeline. Next, ASVs were aligned to the SILVA database^49^ for the taxonomic assignment using the naïve Bayesian classifier implemented by DADA2. Next, raw reads were mapped to the inferred ASVs to generate an abundance table, which was subsequently employed for analyses of diversity and differential abundance using the R package *vegan*^50^.

Amplicon sequencing reads from the *Lotus* and *Arabidopsis*^8^ IRLs and from their corresponding culture-independent root community profiling were quality-filtered and demultiplexed according to their two-barcode (well and plate) identifiers using custom scripts and a combination of tools included in the QIIME^47^ and USEARCH^51^ pipelines. Next, sequences were clustered into Operational Taxonomic Units (OTUs) with a 97% sequence identity similarity using the UPARSE algorithm, followed by identification of chimeras using UCHIME^52^. Samples (wells) with fewer than 100 good quality reads were removed from the data set as well as OTUs not found in a well with at least ten reads. A purity threshold of 90% was chosen for identification of recoverable OTUs. We identified *Lj*-IRL samples matching OTUs found in the culture-independent root samples and selected a set of 294 representative strains maximizing taxonomic coverage for subsequent validation and whole-genome sequencing, forming the basis of the core *Lj*-SPHERE collection.

Sequencing data from SynCom experiments was pre-processed similarly as natural community *16S* data. Quality-filtered, merged paired-end reads were then aligned to a reference set of sequences extracted from the whole-genome assemblies of every strain included in a given gnotobiotic experiment, using USEARCH (*uparse_ref* command)^53^. Only sequences with a perfect match to the reference database were retained. We then checked that the fraction of unmapped reads did not significantly differ between compartment, experiment or host species. Next, we generated a count table that was employed for downstream analyses of diversity with the R package *vegan*^50^. Finally, we visualized amplicon data from all experimental systems using the *ggplot2* R package^54^.

### Host preference and invasiveness indices

In order to quantify the strength of the host preference of each bacterial strain individually, we calculated the ratio between the mean relative abundance of a given SynCom member in root samples of their native host and its mean relative abundance in root samples of the other plant species. To avoid obtaining very high ratios due to small denominator values, strains with mean relative abundances below 0.1% in either of the two hosts were removed from the analysis. Similarly, an invasiveness index was calculated by obtaining the ratio between mean relative abundance of a strain when invading resident communities on roots of their native host, compared to the other plant species. To test whether a SynCom member was significantly more abundant in the roots of their native host (i.e. significant host preference), we used the non-parametric Wilcoxon test controlling for false discovery rate (FDR) with α = 0.05.

### Bacterial genome assembly and annotation

Paired-end Illumina reads were first subjected to length trimming and quality-filtering using Trimmomatic^55^. Subsequently, reads were assembled using the A5 assembly pipeline^56^, which uses the IDBA algorithm^57^ to assemble error-corrected reads. Detailed assembly statistics and corresponding metadata can be found in Supplementary Data 3. Genomes with multi-modal *k*-mer and GC content distributions or multiple instances of marker genes from diverse taxonomic groups were flagged as not originating from clonal cultures. These samples were then processed using a metagenome binning approach^58^. Briefly, contigs from each metagenome sample were clustered using METABAT2^59^, followed by an assessment of completeness and contamination of each metagenome-assembled genome (MAG) using CheckM^60^. Only bins with completeness scores larger than 75% and contamination rates lower than 5% were retained and added to the collection (Supplementary Data 3; designated MAG in the column ‘type’). Next, functional annotation of genes was conducted using Prokka and employing a custom database based on KEGG Orthologue (KO) groups^61^ downloaded from the KEGG FTP server in November 2019. Hits to sequences in the database were filtered using an *E* value threshold of 10 × 10^−9^ and a minimum coverage of 80% of the length of the query sequence.

### Phylogenomic analysis of the *Lj*- and *At*-SPHERE culture collections

Genomes from the *Lj*- and *At*-SPHERE culture collections^8^ were searched for the presence of a set of 31 conserved, single-copy marker genes, known as AMPHORA^62^ genes. Next, sequences of each gene were aligned using Clustal Omega^63^ with default parameters. Using a concatenated alignment of each gene, we inferred a maximum likelihood phylogeny using FastTree^64^. We visualized this tree using the Interactive Tree of Life web tool^65^. Finally, genomes from both collections (*Lj*-SPHERE and *At*-SPHERE) were clustered into phylogroups, roughly corresponding to a species designation^66^ using FastANI^67^ and a threshold of average nucleotide identity at the whole genome level of at least 97%.

### RNA-sequencing and data analysis

RNA isolated from FlowPot samples was subjected to quality control, library preparation, and sequencing (on the Illumina HiSeq3000 platform) at the Max Planck-Genome center, Cologne, Germany (https://mpgc.mpipz.mpg.de/home/). Sequencing depth was 6 million reads persample.

Raw Illumina RNA-Seq reads were pre-processed using fastp (v0.19.10)^68^ with default settings for pair-end reads High quality reads were pseudo-aligned to the *Lotus japonicus* Gifu *Arabidopsis thaliana* Col-0 transcriptome reference using kallisto (v0.46.1)^69^. After removal of low abundant transcripts that were not present in at least two replicates under each condition, count data were imported using the *tximport* package^70^.

Differential expression analyses were performed using the *DESeq2* package^71^. Firstly, raw counts were normalized with respect to the library size (*rlog* function) and transformed into log_2_ scale. We tested for sample effects by surrogate variable (SV) analysis using the *sva* package^72^. Significant SVs were automatically detected and integrated into the model for differential analyses. Principal component analysis based on whole transcripts were then conducted and plotted to visualize the cluster and variance of biological replicates under each condition. Transcripts with fold-changes > 1.5 and adjusted *p*-value for multiple comparisons (Benjamini–Hochberg method) equal to or below 0.05 were considered significant.

The log_2_ scaled counts were normalized by the identified SVs using the *limma* package^73^ (‘removeBatchEffect’ function), and transformed as median-centered *z*-score (by transcripts, ‘scale’ function). Then *z*-scores was used to conduct *k*-means clustering for all transcripts. The cluster number (*k* = 10) was determined by sum of squared error and Akaike information criterion. Differential expressed transcripts and cluster results were visualized using heatmaps generated by *ComplexHeatmap* package^74^.

Gene ontology (GO) enrichment for each cluster using the whole *Lotus* and *Arabidopsis* transcriptomes as backgrounds were performed with the *goseq* package^75^, which considers the transcripts length bias in RNA-Seq data. GO annotations were retrieved from the Gene Ontology Consortium (September 2019)^76,77^. Significantly changed biological process GO terms (adjusted *p*-value < 0.05) were visualized in dot plots using the *clusterProfiler* package^78^.

### Data and code availability

Raw *16S* rRNA amplicon reads will be deposited in the European Nucleotide Archive (ENA) under the accession number PRJEB37695. Similarly, sequencing reads and genome assemblies of the *Lj*-SPHERE core collection will be uploaded to the same database with the accession number PRJEB37696. The scripts used for the computational analyses described in this study are available at http://www.github.com/garridoo/ljsphere, to ensure replicability and reproducibility of these results.

## Acknowledgments

We would like to acknowledge Dr. Paloma Duran for her assistance while performing the SynCom experiments, Anna Lisa Roth and Zuzana Blahovska for their help in maintaining the culture collection, Dr. Jairo Garnica and Dr. Elisa Brambilla for their help in optimizing millifluidics protocols, Dr. Manuela Peukert and Dr. Stanislav Kopriva for their advice on root exudate collection, Dr. Juliette de Meaux for providing *A. lyrata* seeds, and Neysan Donnelly for scientific English editing. This research was funded by the Max Planck Society and Deutsche Forschungsgemeinschaft (DFG, German Research Foundation) under Germany’s Excellence Strategy – EXC-Nummer 2048/1– project 390686111 and the ‘2125 DECRyPT’ Priority Programme through P. S.-L. and R. G.-O. K.T. was funded by Chinese Scholarship Council (CSC). The Novo Nordisk program InRoot, grant number: NNF19SA0059362, funded K.T. and S. R.

## Author contributions

K.W., K. T., S. R., P. S.-L., and R. G.-O. conceived the research and designed the experiments. K. W, K. T., R. Z., and D. B. J. established the *Lj*-SPHERE culture collection. K. W. and K. T. performed the gnotobiotic competition experiments. K. W. and E. L. conducted the *in planta* invasion and millifluidics SynCom experiments. R. G., and R. G.-O. analyzed culture independent amplicon data. E. D., and R. G.-O. analyzed the *Lj*-IRL data. P. Z., and R. G.-O. processed bacterial whole-genome data from the *Lj*-SPHERE collection. Y. N. and R. G.-O. analyzed the transcriptome data. K. W., and R. G.-O. analyzed sequencing data from the SynCom experiments. K. W., K. T., S. R., P. S.-L., and R. G.-O. interpreted data and wrote the paper.

