## Supplementary Material for "Host preference and invasiveness of commensals in the *Lotus* and *Arabidopsis* root microbiota"

**This PDF file includes the following Supplementary Information:**

Supplementary Text
Supplementary Figures 1 to 13
Supplementary Tables 1 and 2

**Supplementary Data guide:**

Supplementary Data 1: Data and metadata of natural community profiles of *Lotus* and *Arabidopsis* roots.

http://www.at-sphere.com/ljsphere/Supplementary_Data_S1.xlsx

Supplementary Data 2: Data and metadata of *Lj*- and *At*-IRLs.

http://www.at-sphere.com/ljsphere/Supplementary_Data_S2.xlsx

Supplementary Data 3: Metadata of the *Lj*- and *At*-SPHERE core culture collection.

http://www.at-sphere.com/ljsphere/Supplementary_Data_S3.xlsx

Supplementary Data 4: Data and metadata of *LjAt* SynCom experiments.

http://www.at-sphere.com/ljsphere/Supplementary_Data_S4.xlsx

**Supplementary Text**

Culture collection recovery rates

To explore the mechanisms by which different plant species assemble distinct microbial communities, we established a taxonomically and functionally diverse culture collection of the *Lotus* root and nodule microbiota (Methods). A total of 3,960 colony-forming units (CFUs) were obtained and taxonomically characterized by sequencing the bacterial *16S* ribosomal RNA (rRNA; Supplementary Data 2), resulting in a comprehensive sequence-indexed rhizobacterial library from *L. japonicus* (*Lj*-IRL). In parallel, a subset of the root samples was also subjected to amplicon sequencing to obtain culture-independent community profiles for cross-referencing with the *Lj*-IRL data. Recovery rates were estimated by calculating the number of bacterial OTUs (Operational Taxonomic Units, defined by 97% sequence identity) found in the natural communities that had at least one isolate in our culture collection (Methods). For *Lotus*, the recovery rates varied between 50% (based on the top 100 most abundant OTUs), 53% (OTUs with RA $\geq$ 0.1%), and 64.58% (prevalent OTUs, found in at least 80% of the natural community samples). Recovered OTUs accounted for up to 82% of the cumulative relative abundance of the entire culture-independent community (Fig. 1c), indicating that our collection is representative of a large fraction of the *Lotus* root microbiota. By comparison, the recovery rates for the *A. thaliana* culture collection (*At*-IRL) varied between 51% (top 100 OTUs), 57% ($\geq$ 0.1% relative abundance), and 62.82% (prevalent OTUs), while recovered OTUs recovered from *Arabidopsis* roots reached a cumulative relative abundance of 59% of the entire community (Fig. 1e). Interestingly, 45.57% of the abundant OTUs found in the natural communities of *Lj* roots were recovered in the *At*-IRL, whereas 45.19% of abundant OTUs from *At* roots were recovered in the *Lj*-IRL (Fig. 1d, and 1f). These results are indicative of a substantial overlap of the recovered bacterial OTUs.

Taxonomic and functional overlap of the *Lotus* and *Arabidopsis* culture collections

To establish a core *Lotus* culture collection of whole-genome sequenced strains (*Lj*-SPHERE), we selected from the *Lj*-IRL a taxonomically representative subset of bacterial isolates maximizing the number of taxa covered (Methods). A total of 294 isolates belonging to 20 families and 124 species, including both commensal and symbiotic bacteria, were subjected to whole-genome sequencing (Supplementary Data 3). This core collection is of a similar size and diversity as the collection from *Arabidopsis* roots (*At*-SPHERE)[8]. A whole-genome phylogeny of all sequenced isolates from both collections revealed an extensive taxonomic overlap between exemplars derived from *Lotus* and *Arabidopsis* (Supplementary Fig. 2), indicating that the observed differences in natural community structures (Fig. 1b) are likely not driven by the presence of host-specific bacterial taxonomic groups. Instead, the distinct root community profiles of the two hosts are possibly due to differences in the relative abundance of shared taxonomic groups (Supplementary Fig. 1).

We hypothesized that bacterial preference for a plant species should be accompanied by the acquisition of a set of genes required for preferential colonization of a specific host. In order to test this, we characterized the functional potential encoded in the genomes of the sequenced isolates using the KEGG orthology database as a reference[61]. We observed that a large proportion of annotated gene families was shared between the two culture collections (6,712 out of 7,456), and that the number of gene families exclusively found in genomes of strains derived from *Lotus* or *Arabidopsis* roots (3.51% and 6.47%, respectively) did not significantly deviate from what would be expected by chance ($P$ = 0.49). However, additional host-specific genes are likely encoded in sequences for which a functional annotation is currently unavailable (~27%). Principal coordinates analysis (PCoA) of functional distances revealed a high degree of overlap between isolates of the same taxonomic groups, which was independent of their host of origin (Supplementary Fig. 2). Permutation analysis of variance confirmed that the main driver of functional variation encoded in the genomes of our culture collections was the taxonomy of the isolates (79.70% of variance explained; $P$ = 0.001), and that the origin of isolation (i.e., host species) only explained a small fraction of the functional diversity encoded by these genomes (4.27% of variance; $P$ = 0.001).

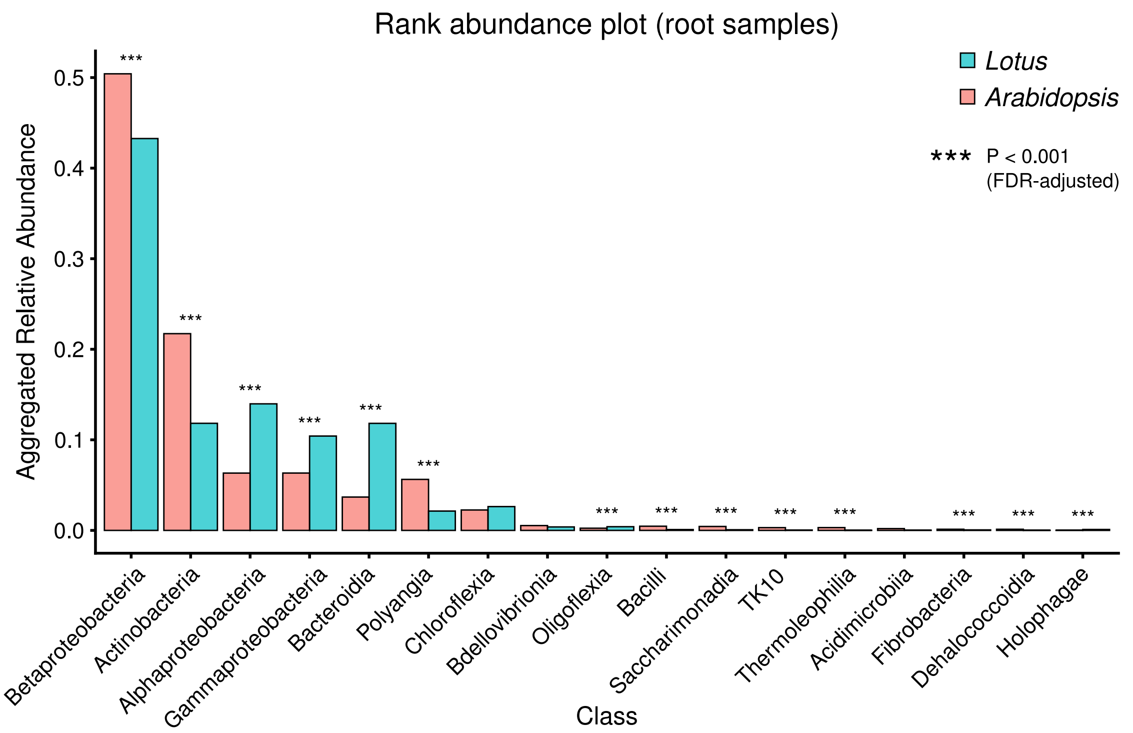


**Supplementary Figure 1 | Culture-independent diversity analysis of root-associated bacterial communities from *Lotus* and *Arabidopsis*.** Rank abundance plot of bacterial communities from *Lotus* or *Arabidopsis* roots, aggregated to the class level, including all families with a mean accumulated relative abundance of > 0.1% on either host.

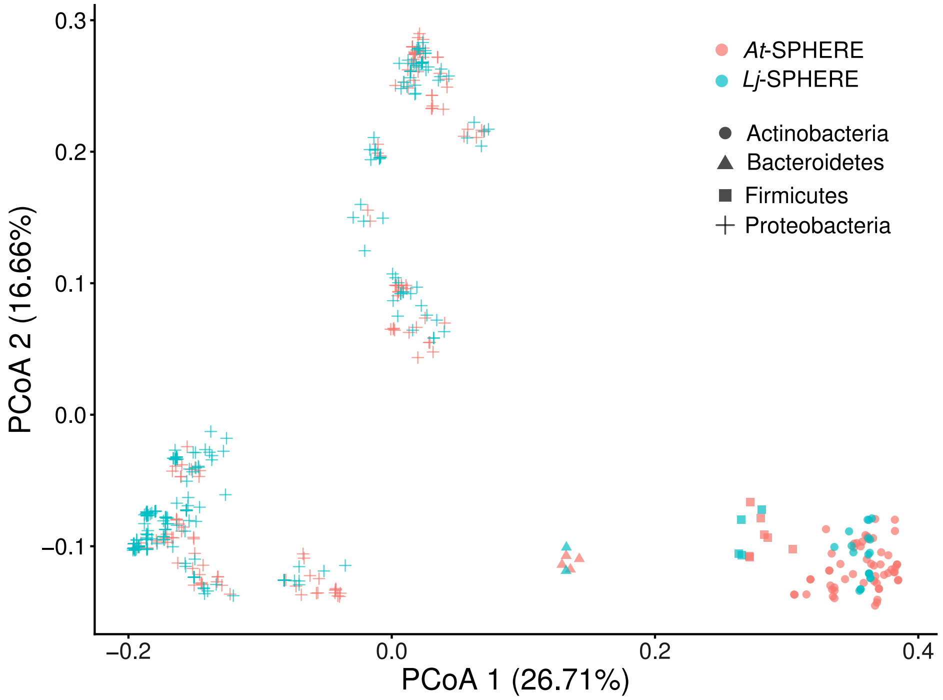


**Supplementary Figure 2 | Functional overlap of *Lotus* and *Arabidopsis* culture collection genomes.** PCoA of functional distances of genomes from bacterial isolates of the *Lotus* (*Lj*-SPHERE; $n$ = 294) and the *Arabidopsis* (*At*-SPHERE; $n$ = 194) culture collections.

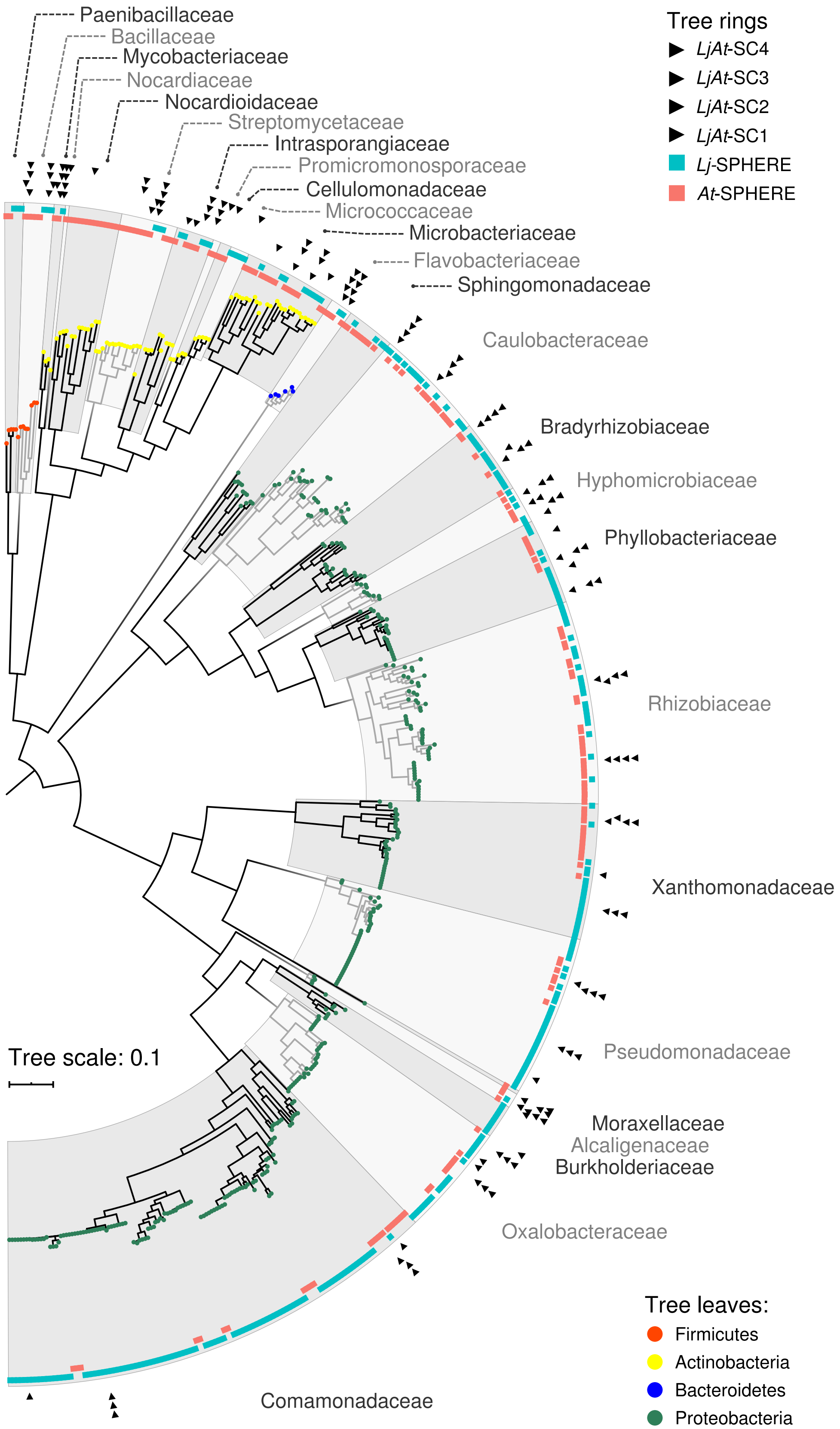
Tree rings
LjAt-SC4
LjAt-SC3
LjAt-SC2
LjAt-SC1
Lj-SPHERE
At-SPHERE
Paenibacillaceae
Bacillaceae
Mycobacteriaceae
Nocardiaceae
Nocardioidaceae
Streptomycetaceae
Intrasporangiaceae
Promicromonosporaceae
Cellulomonadaceae
Micrococcaceae
Microbacteriaceae
Flavobacteriaceae
Sphingomonadaceae
Caulobacteraceae
Bradyrhizobiaceae
Hyphomicrobiaceae
Phyllobacteriaceae
Rhizobiaceae
Xanthomonadaceae
Pseudomonadaceae
Moraxellaceae
Alcaligenaceae
Burkholderiaceae
Oxalobacteraceae
Comamonadaceae
Tree scale: 0.1
Tree leaves:
Firmicutes
Actinobacteria
Bacteroidetes
Proteobacteria

**Supplementary Figure 3 | Whole-genome phylogeny of the *Lotus* and *Arabidopsis* core culture collections.** Maximum likelihood phylogeny, constructed from a concatenated alignment of 31 conserved, single copy genes (AMPHORA) showing the taxonomic overlap of the *Lj*-SPHERE (*n* = 294, blue track) and *At*-SPHERE (*n* = 194, red track) core culture collections. Arrows in the outer rings indicate the strains selected for four mixed communities used in reconstitution experiments.

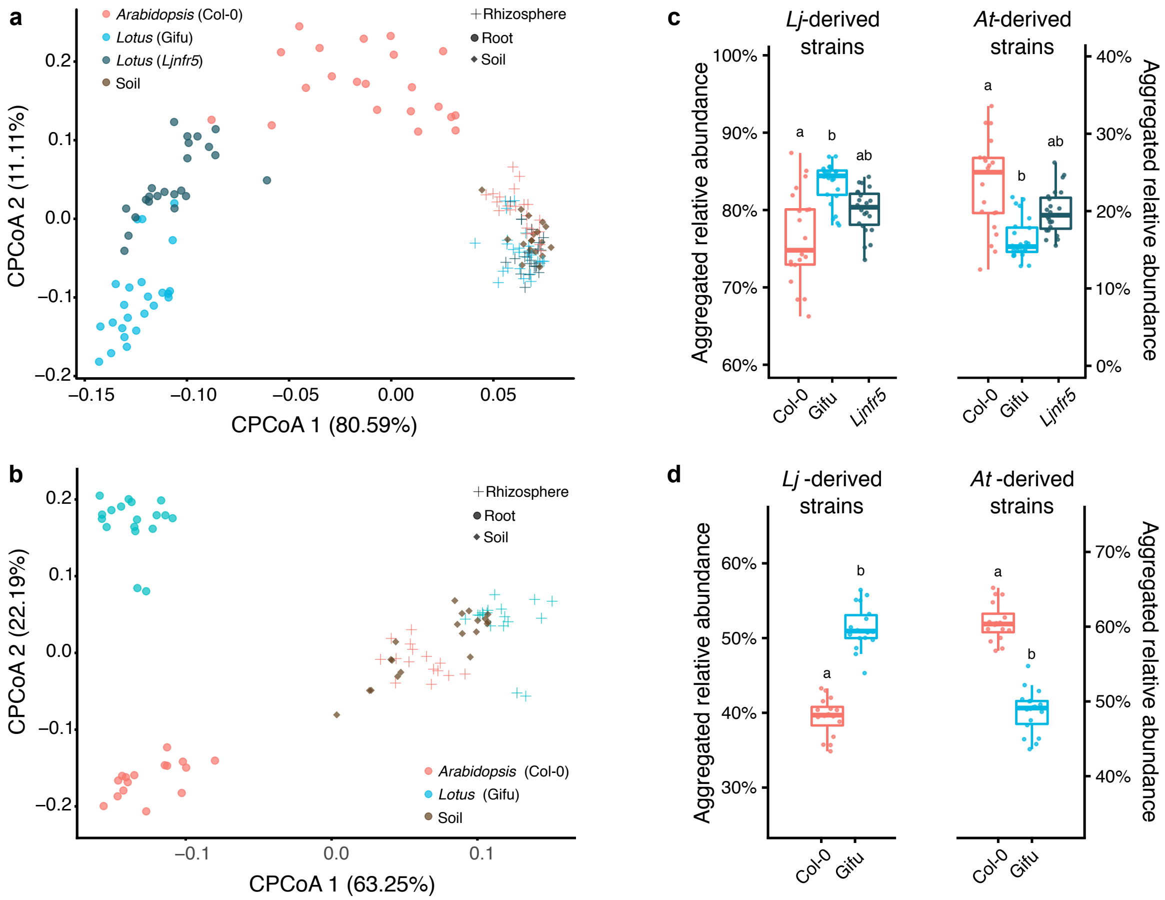


**Supplementary Figure 4 | Host-species specific bacterial root communities and commensal host preference is confirmed using independent mixed communities. a** and **c**, Constrained PCoA of Bray-Curtis dissimilarity (constrained by all biological factors and conditioned by all technical variables) of soil, rhizosphere, and root samples from *L. japonicus* wild type Gifu, *nfr5* mutant, and *A. thaliana* wild type Col-0 plants co-cultivated with the mixed community *LjAt*-SC1 (**a**, experiment A, $n$ = 155, variance explained 53.8%, $P$ = 0.001), or from Gifu and Col-0 co-cultivated with *LjAt*-SC4 (**b**, experiment C, $n$ = 87, variance explained 60%, $P$ = 0.001). **c** and **d**, Aggregated RA of the 16 *Lj*-derived and the 16 *At*-derived strains in the roots of *Lotus* and *Arabidopsis* plants inoculated with *LjAt*-SC1 (**c;** $n$ = 68), or *LjAt*-SC4 (**d**; $n$ = 34).

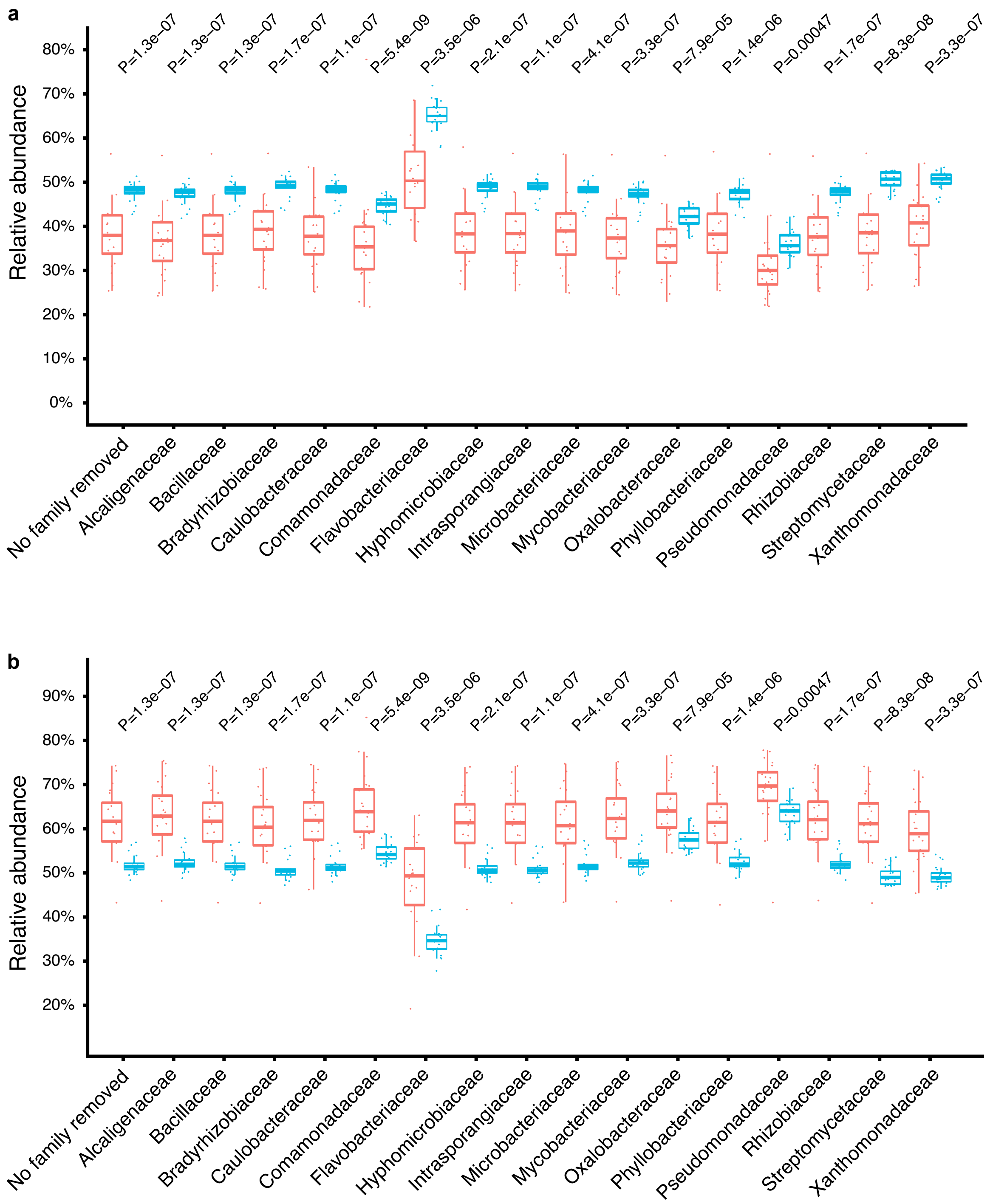


**Supplementary Figure 5 | Host preference is retained after *in silico* removal of individual bacterial families.** Aggregated relative abundance of *Lotus*- (**a**) and *Arabidopsis*-derived (**b**) strains in roots from plants inoculated with the mixed community *LjAt*-SC3. Host preference was assessed using a Mann-Whitney non-parametric test after *in silico* removal of each family. The x-axis labels indicate each depleted family.

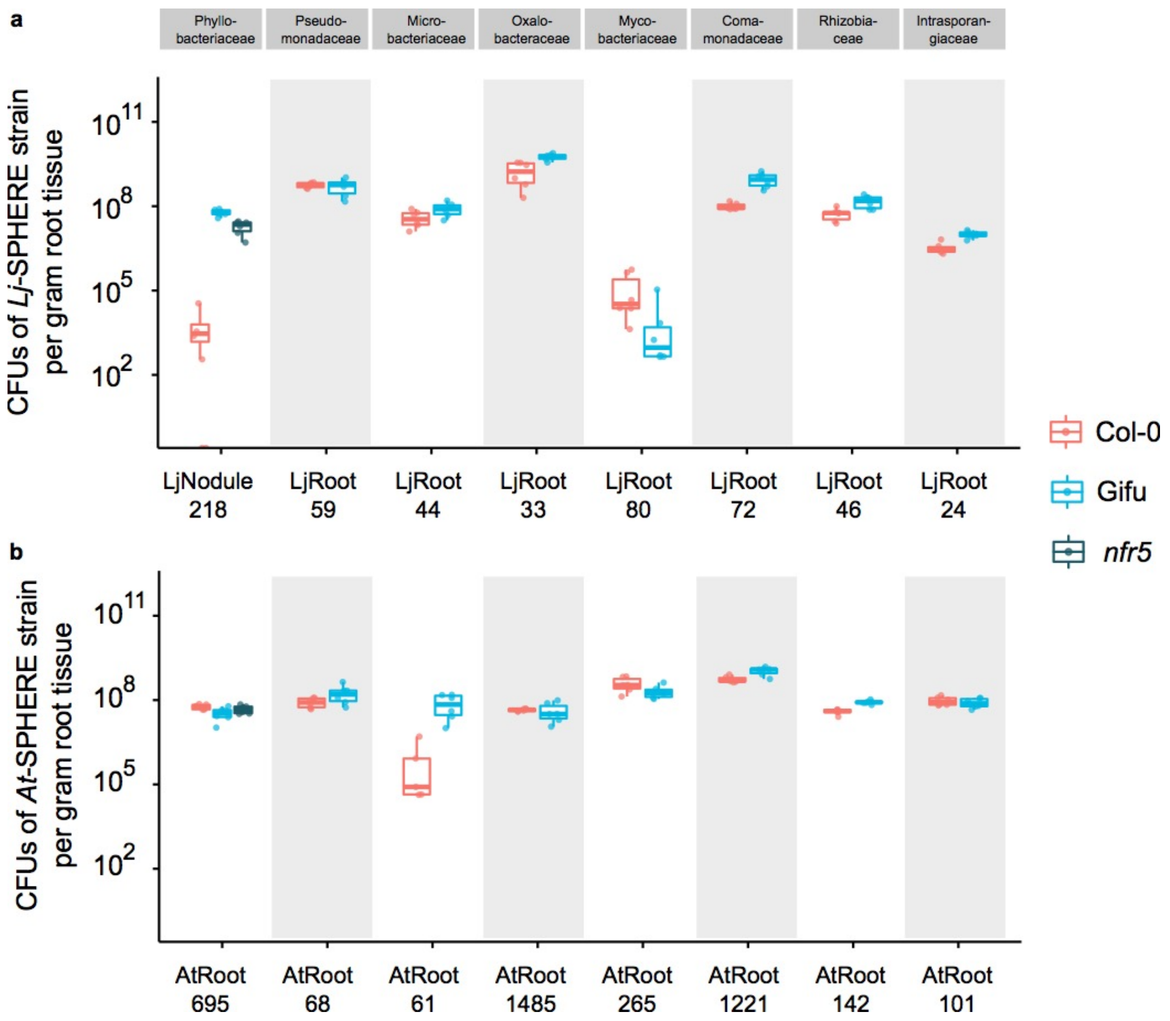


**Supplementary Figure 6 | Bacterial abundance in mono-association with host plants.** Bacterial abundances of commensal bacteria colonizing Col-0, Gifu, and *nfr5* roots, assessed by counting of colony forming units (CFUs) after extraction from root tissue. Plants were grown for two weeks on agar plates in mono-association with the indicated *Lj*-SPHERE (**a**) and *At*-SPHERE (**b**) strains.

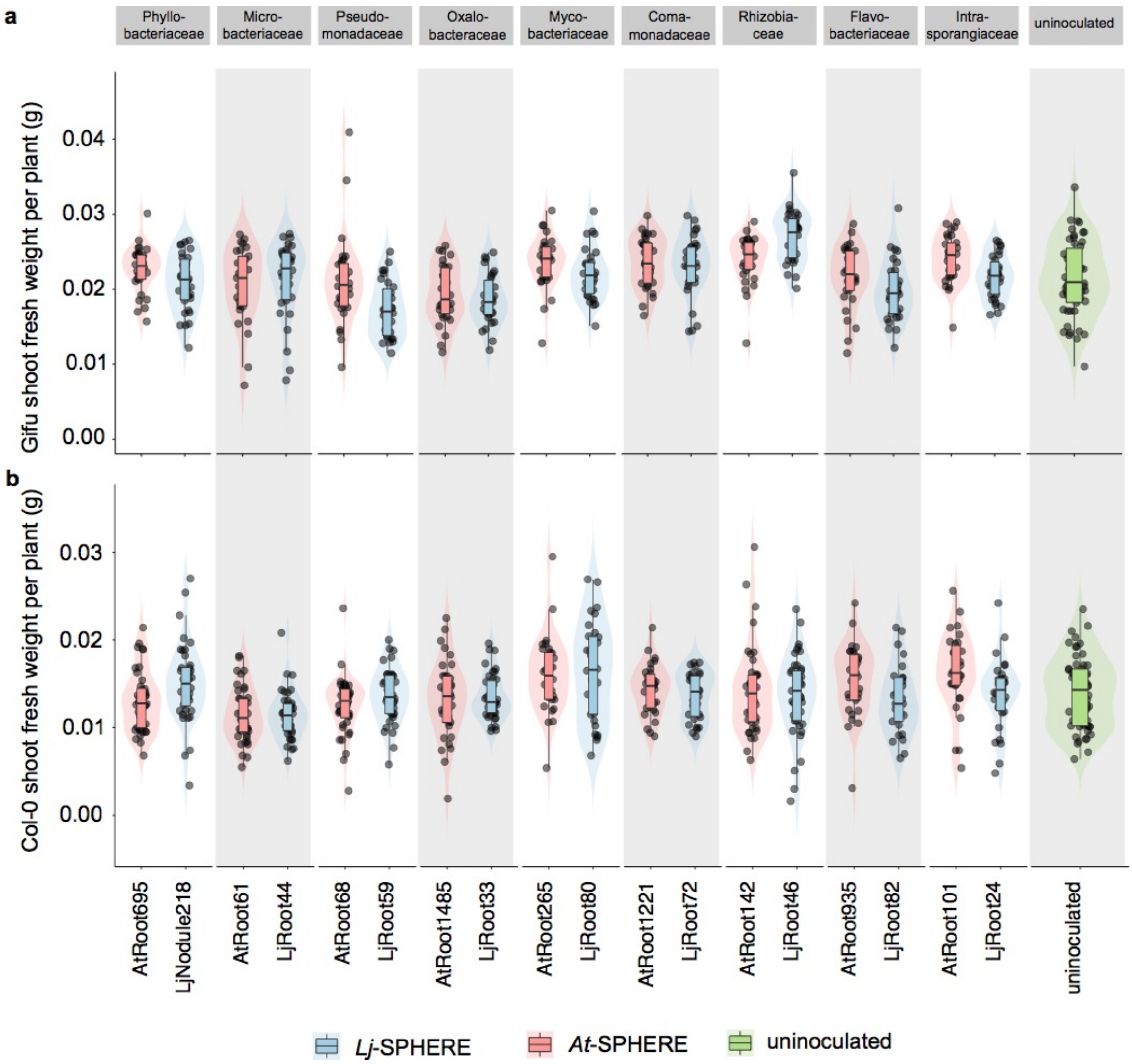


**Supplementary Figure 7 | Plant performance in mono-associations.** Shoot fresh weight of Gifu (**a**) and Col-0 (**b**) plants grown for two weeks on agar plates in mono-association with the indicated *Lj*-SPHERE and *At*-SPHERE strains.

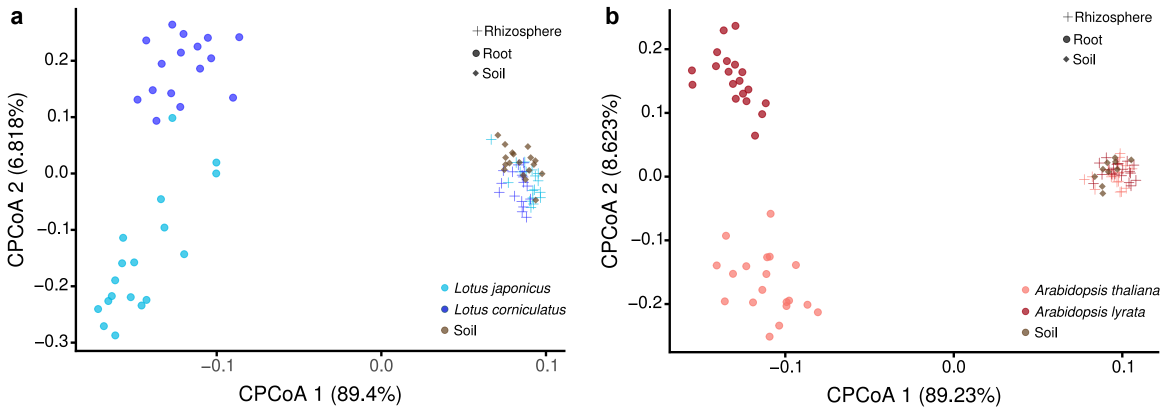


**Supplementary Figure 8 | Sister species of *L. japonicus* and *A. thaliana* establish distinct bacterial root communities. a** and **b**, Constrained PCoA of Bray-Curtis dissimilarity (constrained by all biological factors and conditioned by all technical variables) of root samples from *L. japonicus* wild type Gifu and *L. corniculatus* (**a**; $n$ = 87; variance explained 58.7%, $P$=0.001), and or root samples from *A. thaliana* wild type Col-0 and *A. lyrata* MN47 (**b**; $n$ = 86; variance explained 65%, $P$=0.001), inoculated and grown with the mixed community *LjAt*-SC3, and of the corresponding rhizosphere and bulk soil communities.

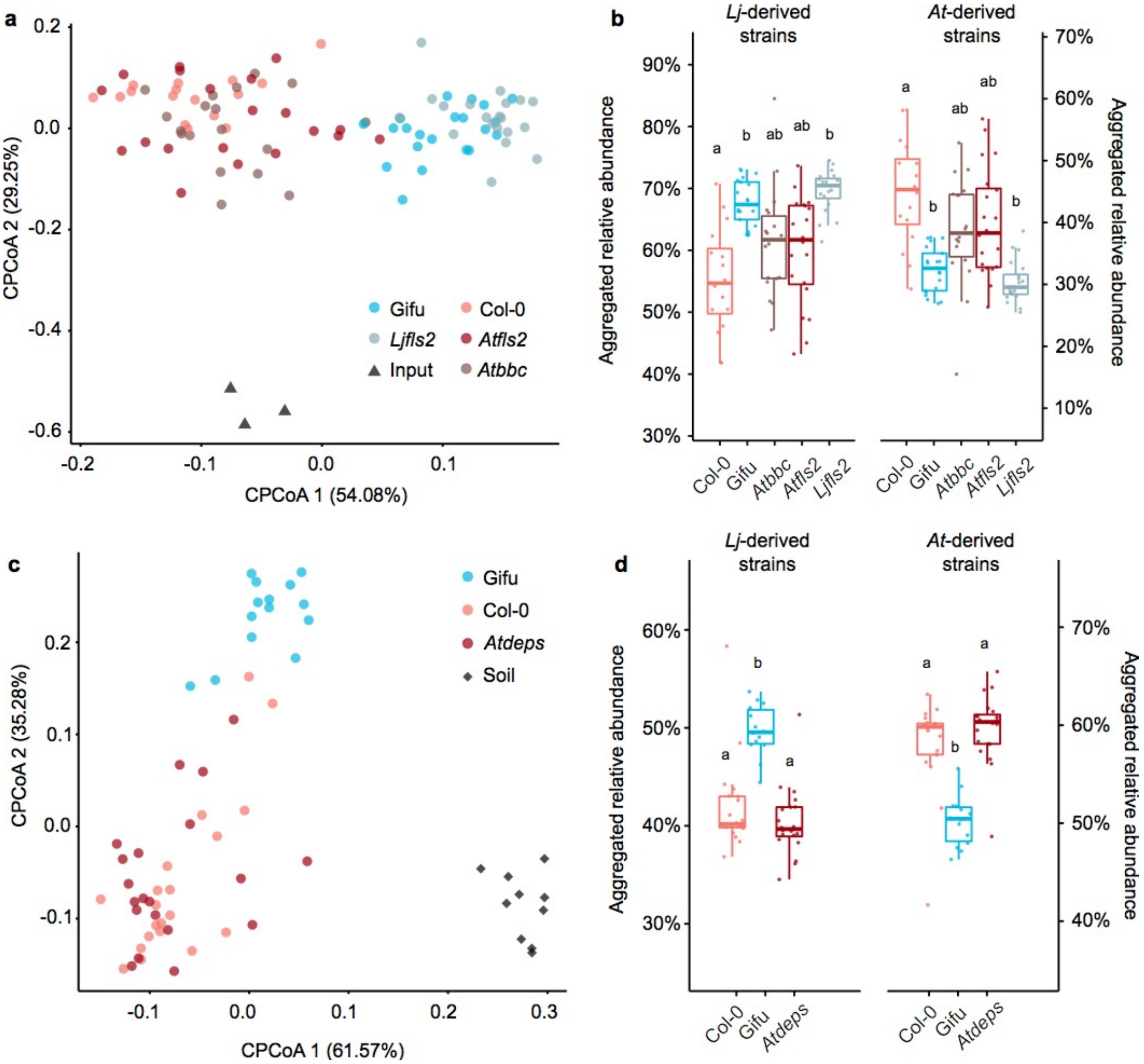


**Supplementary Figure 9 │ Tested plant immune receptors and signaling pathways do not affect host preference of commensals. a**, Constrained PCoA of Bray-Curtis dissimilarity (constrained by all biological factors and conditioned by all technical variables; $n$ = 98; variance explained 24.6%, $P$=0.001) of root samples from *L. japonicus* wild type Gifu, *Ljfls2* mutant, *A. thaliana* wild type Col-0, *Atfls2* mutant, and *Atbbc* mutant inoculated and grown with the mixed SynCom *LjAt*-SC1, and of the corresponding bacterial input communities. **b**, Aggregated relative abundance of the 16 *Lj*-derived and the 16 *At*-derived strains in the roots of *Lotus* and *Arabidopsis* plants. **c**, Constrained PCoA of Bray-Curtis dissimilarity (constrained by all biological factors and conditioned by all technical variables; $n$ = 64; variance explained 42.2%, $P$=0.001) of soil and root samples from Gifu, Col-0, and *Atdeps* mutant inoculated and grown with the mixed SynCom *LjAt*-SC3. **d**, Aggregated relative abundance of the 16 *Lj*-derived and the 16 *At*-derived strains in the roots of *Lotus* and *Arabidopsis* plants.

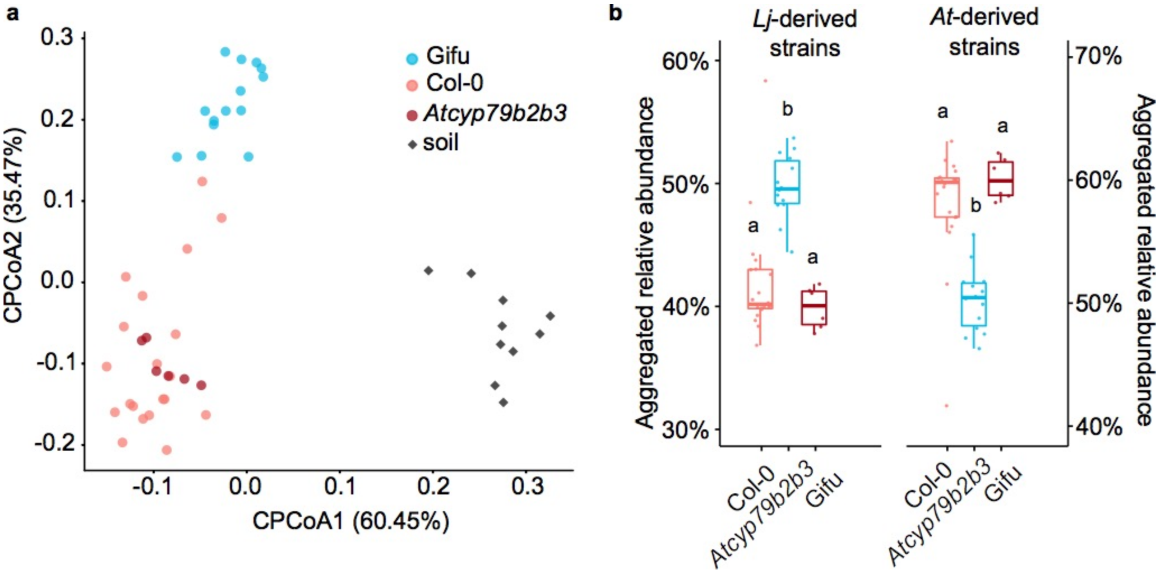


**Supplementary Figure 10 │ Effect of secreted indole glucosinolates on host preference of commensals. a**, Constrained PCoA of Bray-Curtis dissimilarity (constrained by all biological factors and conditioned by all technical variables; *n* = 50; variance explained 47.2%, *P*=0.001) of soil and root samples from *L. japonicus* wild type Gifu, *A. thaliana* wild type Col-0, and *Arabidopsis cyp79b2 cyb79b3* mutant inoculated and grown with the mixed SynCom *LjAt*-SC3. **b**, Aggregated relative abundance of the 16 *Lj*-derived and the 16 *At*-derived strains in the roots of *Lotus* and *Arabidopsis* plants.

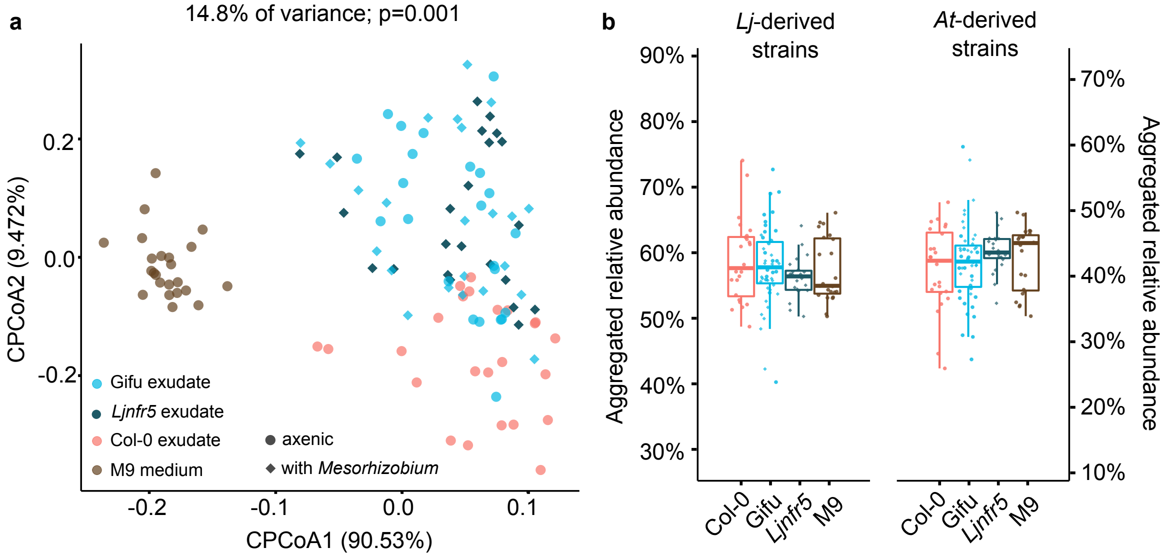


**Supplementary Figure 11 │ Effect of soluble root exudates on host preference of commensals. a**, Constrained PCoA of Bray-Curtis dissimilarity (constrained by all biological factors and conditioned by all technical variables; $n = 116$) of the mixed SynCom *LjAt*-SC1 incubated in root exudates from axenically grown Gifu or Col-0, from Gifu or *Ljnfr5* inoculated with the symbiont *Mesorhizobium*, or in a carbon-rich control medium M9. **b**, Aggregated relative abundance of the 16 *Lj*-derived and the 16 *At*-derived strains in the *Lotus* and *Arabidopsis* exudates. A Kruskal-Wallis test showed no significant differences in the distribution of values among groups.

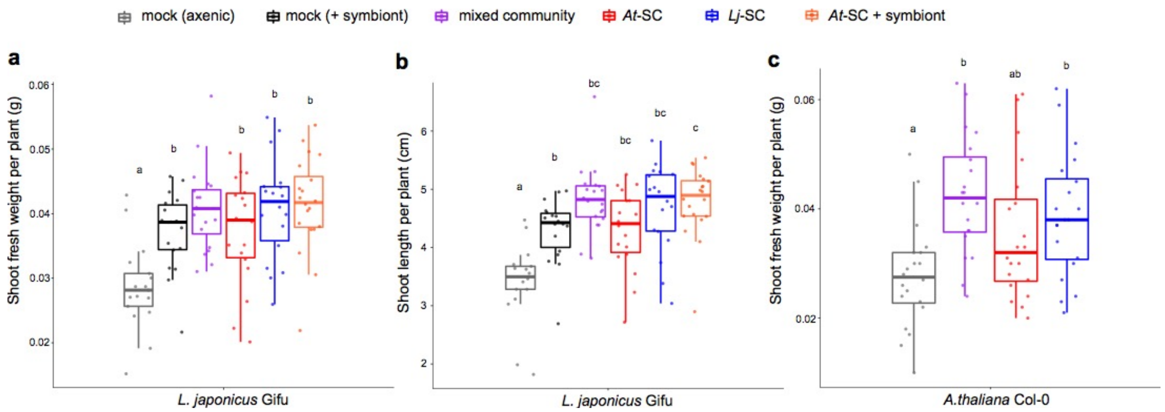


**Supplementary Figure 12 | Shoot phenotypes of *Lotus* and *Arabidopsis* plants inoculated with different commensal communities.** *L. japonicus* Gifu and *A. thaliana* Col-0 plants were co-cultivated with the mixed community *LjAt*-SC3, or individual SynComs *Lj*-SC3 and *At*-SC3. Shoot fresh weight of *Lotus* (**a**) and *Arabidopsis* (**c**), as well as shoot length of *Lotus* (**b**) were measured after five weeks. Each data point corresponds to one replicate comprising roots of 2-4 plants grown in the same pot. Shared letters indicate no significant difference based on Kruskal-Wallis and Wilcoxon rank sum test ($P < 0.05$, $n$ = 18-20).

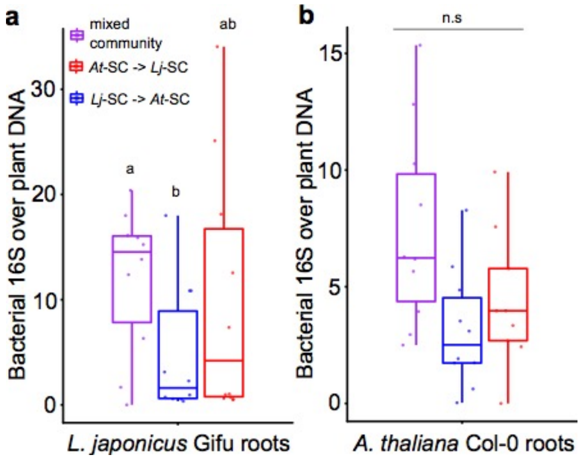


**Supplementary Figure 13 | Quantification of bacterial load on plant roots after sequential inoculation with native and non-native commensals.** *L. japonicus* Gifu and *A. thaliana* Col-0 plants were co-cultivated with the mixed community *LjAt*-SC3, or individual SynComs *Lj*-SC3 and *At*-SC3, followed by inoculation with the remaining SynCom. Colors relate to the early-arriving community. Amount of 16S rRNA gene copies relative to plant gene copies as proxy for bacterial load on *Lotus* (**a**) and *Arabidopsis* (**b**) roots is shown. Each data point corresponds to one replicate comprising roots of 2-4 plants grown in the same pot. Shared letters indicate no significant difference based on Kruskal-Wallis and Wilcoxon rank sum test ($P < 0.05$, $n = 10$-14).

**Supplementary Table 1 | Bacterial SynComs used in this study**

| | | *LjAt*-SC1 | | *LjAt*-SC2 | | *LjAt*-SC3 | | *LjAt*-SC4 | |
|---|---|---|---|---|---|---|---|---|---|
| Class | Family | *At*-SC1 | *Lj*-SC1 | *At*-SC2 | *Lj*-SC2 | *At*-SC3 | *Lj*-SC3 | *At*-SC4 | *Lj*-SC4 |
| Betaproteobacteria | Alcaligenaceae | AtRoot83 | LjRoot1 | AtRoot170 | LjRoot1 | AtRoot83 | LjRoot1 | AtRoot83 | LjRoot1 |
| Firmicutes | Bacillaceae | AtRoot131 | LjRoot5 | AtRoot11 | LjRoot53 | AtRoot131 | LjRoot5 | AtRoot131 | LjRoot5 |
| Alphaproteobacteria | Bradyrhizobiaceae | AtRoot123D2 | LjRoot52 | AtRoot123D2 | LjRoot4 | AtRoot123D2 | LjRoot52 | AtRoot123D2 | LjRoot52 |
| Alphaproteobacteria | Caulobacteraceae | AtRoot77 | LjRoot17 | AtRoot1290 | LjRoot17 | AtRoot77 | LjRoot17 | AtRoot77 | LjRoot17 |
| Betaproteobacteria | Comamonadaceae | AtRoot404 | LjRoot109 | AtRoot16D2 | LjRoot72 | AtRoot1221 | LjRoot72 | AtRoot1221 | LjRoot72 |
| Bacteroidetes | Flavobacteriaceae | AtRoot186 | LjRoot149 | AtRoot935 | LjRoot82 | AtRoot935 | LjRoot82 | AtRoot935 | LjRoot82 |
| Alphaproteobacteria | Hyphomicrobiaceae | AtRoot436 | LjRoot16 | AtRoot436 | LjRoot3 | AtRoot685 | LjRoot16 | AtRoot685 | LjRoot16 |
| Actinobacteria | Intrasporangiaceae | AtRoot85 | LjRoot27 | AtRoot101 | LjRoot24 | AtRoot101 | LjRoot24 | AtRoot101 | LjRoot24 |
| Actinobacteria | Microbacteriaceae | AtRoot61 | LjRoot44 | AtRoot4 | LjRoot42 | AtRoot61 | LjRoot44 | AtRoot61 | LjRoot44 |
| Actinobacteria | Mycobacteriaceae | AtRoot265 | LjRoot80 | AtRoot135 | LjRoot80 | AtRoot265 | LjRoot80 | AtRoot265 | LjRoot80 |
| Betaproteobacteria | Oxalobacteraceae | AtRoot335 | LjRoot35 | AtRoot1485 | LjRoot33 | AtRoot1485 | LjRoot33 | AtRoot1485 | LjRoot33 |
| Alphaproteobacteria | Phyllobacteriaceae | AtRoot695 | LjNodule218 | AtRoot554 | LjNodule210 | AtRoot695 | LjNodule218 | AtRoot695 | LjNodule218 |
| Gammaproteobacteria | Pseudomonadaceae | AtRoot71 | LjRoot54 | AtRoot68 | LjRoot59 | AtRoot68 | LjRoot59 | AtRoot68 | LjRoot59 |
| Alphaproteobacteria | Rhizobiaceae | AtRoot142 | LjRoot46 | AtRoot142 | LjRoot2 | AtRoot142 | LjRoot46 | AtRoot142 | LjRoot46 |
| Actinobacteria | Streptomycetaceae | AtRoot63 | LjRoot303 | AtRoot1295 | LjRoot303 | AtRoot1310 | LjRoot303 | AtRoot1310 | LjRoot303 |
| Gammaproteobacteria | Xanthomonadaceae | AtRoot480 | LjRoot21 | AtRoot627 | LjRoot60 | AtRoot480 | LjRoot60 | AtRoot480 | LjRoot60 |
| Actinobacteria | Cellulomonadaceae | | | | | | | AtRoot137 | |
| Gammaproteobacteria | Moraxellaceae | | | | | | | AtRoot1280 | |
| Actinobacteria | Nocardiaceae | | | | | | | AtRoot136 | |
| Actinobacteria | Nocardioidaceae | | | | | | | AtRoot224 | |
| Actinobacteria | Promicromonosporaceae | | | | | | | AtRoot22 | |
| Betaproteobacteria | Burkholderiaceae | | | | | | | | LjRoot22 |
| Actinobacteria | Micrococcaceae | | | | | | | | LjRoot78 |

| |
|---|
| present in *LjAt*-SC1 |
| present in *LjAt*-SC2 |
| present in all four mixed SynComs |
| present in *LjAt*-SC3 and *LjAt*-SC4 |
| members of host-specific families |

*LjAt*-SC1 was used in experiment A, D, and H (see Supplementary Table 2).

*LjAt*-SC2 was designed for experiment B, (full-factorial replicate of A) to comprise independent strains of the same families, as far as possible (distinguishable 16S sequence).

*LjAt*-SC3 was built using strains from LjAt-SC1 and LjAt-SC2 to generate an idependent community.

*LjAt*-SC4 is identical to *LjAt*SC3, but includes strains from host-specific bacterial families.

In general, mixed communites were designed to include strains distinguishable based on *16S* rRNA gene sequence, to have a similar number of strains in the *Lj* and *At* SynCom, to consist of taxonomically paired *Lj* and *At* SynComs (so that any differences in community structure would be attributable to the origin of strain isolation, i.e., the host plant). In addition, the SynCom design was influenced by practical constraints, e.g., not all strains of the current culture collection or their genome sequences were available at the time of experiment setup, and occasionally strains had to be cured from small contaminations.

**Supplementary Table 1 | Bacterial SynComs used in this study**

**Supplementary Table 2 | List of SynCom experiments**

| ID | Name | Treatments | Second inoculation (after 4 weeks) | Genotypes | Compartments harvested | Number of samples | Growth period |
|---|---|---|---|---|---|---|---|
| A | AtLj_001 | *LjAt*-SC1 | | Gifu | root | 24 | 5 weeks |
| | | | | | rhizosphere | 24 | |
| | | | | Col-0 | root | 21 | |
| | | | | | rhizosphere | 22 | |
| | | | | *Ljnfr5* | root | 23 | |
| | | | | | rhizosphere | 23 | |
| | | | | unplanted | soil | 18 | |
| B | AtLj_002 | *LjAt*-SC2 | | Gifu | root | 24 | 5 weeks |
| | | | | | rhizosphere | 24 | |
| | | | | Col-0 | root | 19 | |
| | | | | | rhizosphere | 24 | |
| | | | | *Ljnfr5* | root | 23 | |
| | | | | | rhizosphere | 24 | |
| | | | | unplanted | soil | 18 | |
| C | AtLj_006 | *LjAt*-SC4 | | Gifu | root | 18 | 5 weeks |
| | | | | | rhizosphere | 18 | |
| | | | | Col-0 | root | 16 | |
| | | | | | rhizosphere | 15 | |
| | | | | unplanted | soil | 20 | |
| D | AtLj 003 | *LjAt*-SC1 | | Gifu | root | 21 | 5 weeks |
| | | | | Col-0 | root | 16 | |
| | | | | *Ljfls2* | root | 20 | |
| | | | | *Atfls2* | root | 20 | |
| | | | | *Atbbc* | root | 18 | |
| E | AtLj 005 | *LjAt*-SC3 | | Gifu | root | 14 | 5 weeks |
| | | | | Col-0 | root | 20 | |
| | | | | *Atdeps* | root | 20 | |
| | | | | *Atcyp79b2 Atcyp79b3* | root | 6 | |
| | | | | unplanted | soil | 10 | |
| F | AtLj_004 | *LjAt*-SC3 | mock | Gifu | root | 20 | 6 weeks |
| | | | | | rhizosphere | 20 | |
| | | | | Col-0 | root | 20 | |
| | | | | | rhizosphere | 20 | |
| | | | | unplanted | soil | 10 | |
| | | *Lj*-SC3 | *At*-SC3 | Gifu | root | 10 | |
| | | | | | rhizosphere | 20 | |
| | | | | Col-0 | root | 20 | |
| | | | | | rhizosphere | 20 | |
| | | | | unplanted | soil | 10 | |
| | | *At*-SC3 | *Lj*-SC3 | Gifu | root | 19 | |
| | | | | | rhizosphere | 19 | |
| | | | | Col-0 | root | 19 | |
| | | | | | rhizosphere | 20 | |
| | | | | unplanted | soil | 10 | |
| G | AtLj_007 | *LjAt*-SC3 | | Gifu | root | 18 | 5 weeks |
| | | | | | rhizosphere | 18 | |
| | | | | Col-0 | root | 19 | |
| | | | | | rhizosphere | 19 | |
| | | | | *L. corniculatus* wild type | root | 16 | |
| | | | | | rhizosphere | 17 | |
| | | | | *A. lyrata* MN47 | root | 19 | |
| | | | | | rhizosphere | 19 | |
| | | | | unplanted | soil | 28 | |
| H | MDA10 | *LjAt*-SC1 | | Gifu (exudates) | droplets | 47 | 3 days |
| | | | | Col-0 (exudates) | droplets | 24 | |
| | | | | *Ljnfr5* (exudates) | droplets | 23 | |
| I | AtLj 008 | *LjAt*-SC3 | | Gifu (dead roots) | dead root | 45 | 5, 12, 19 days |
| | | | | | detritusphere | 45 | |
| | | | | Col-0 (dead roots) | dead root | 44 | |
| | | | | | detritusphere | 45 | |
| | | | | toothpick | wood | 35 | |
| | | | | unplanted | soil | 36 | |

Gifu, *L. japonicus* wild type; Col-0, *A. thaliana* wild type; SC, synthetic community (see also Supplementary Table 1

**Supplementary Table 2 | List of SynCom experiments**
